# PRC domain-containing proteins modulate FtsZ-based archaeal cell division

**DOI:** 10.1101/2023.03.28.534543

**Authors:** Phillip Nußbaum, Danguole Kureisaite-Ciziene, Dom Bellini, Chris van der Does, Marko Kojic, Najwa Taib, Simonetta Gribaldo, Martin Loose, Jan Löwe, Sonja-Verena Albers

## Abstract

Dividing cells into two daughter cells is a complicated process that in bacteria and eukaryotes requires many proteins to work together. For archaea that divide via an FtsZ-based mechanism, only three proteins of the cell division machinery could so far be identified. These are two tubulin homologs, FtsZ1, FtsZ2 and the membrane anchor of FtsZ2, SepF. Here, we investigate additional archaeal cell division proteins that were identified by immunoprecipitation of SepF. These proteins comprise a single PRC-barrel domain and strictly co-occur with FtsZ. Two out of three PRC-barrel domain containing proteins found in *Haloferax volcanii*, CdpB1 and CdpB2 localize to the site of cell division in a SepF-dependent manner. Moreover, depletions and deletions cause severe cell division defects, generating drastically enlarged cells. Fluorescence microscopy of tagged FtsZ1, FtsZ2 and SepF in CdpB1/2 deletion strains revealed that the divisome is unusually disordered and not organized into a distinct ring-like structure at the cell centre. Biochemical analysis of CdpB homologs from different archaeal species showed that SepF interacts directly with CdpB1, which in turn binds to CdpB2, forming a tripartite complex. A crystal structure of CdpB1 and B2 recapitulated these interactions and suggested how these proteins might form filaments, possibly aligning SepF and therefore the FtsZ2 ring during cell division. In summary, we demonstrate that PRC domain proteins play essential roles in FtsZ based cell division in archaea.

## Introduction

Cell division is the fundamental process for cells to generate progeny and to pass on their genetic content to the next generation. In well-studied organisms, this process is known to be executed by many proteins working together to ensure proper division. For example, in bacteria, division is organized by the cell division protein and tubulin homolog FtsZ (1), whereas eukaryotic cells divide using proteins of the ESCRT-system (endosomal sorting complex required for transport) for final abscission (2). Interestingly, in Archaea, both FtsZ-and ESCRT III-based cell division systems exist (3, 4). Current knowledge about the archaeal FtsZ-based cell division system is limited, and only two proteins homologous to bacterial cell division proteins have been characterized recently in archaea (4). One is the GTP-dependent FtsZ (5–8) and the only other is one of the FtsZ membrane anchors found in Firmicutes, Actinobacteria and Cyanobacteria (9, 10), SepF (7, 8). Most archaea that use FtsZ-based cell division contain two phylogenetically distinct FtsZ homologs, including the model organism *Haloferax volcanii* (4). The roles of both FtsZ homologs, FtsZ1 and FtsZ2 during cell division have recently been investigated, showing that they have distinct roles (6). Both tubulin homologs are localized together at the site of cell division, however, FtsZ2 depends on the presence of FtsZ1 for proper localization, whilst FtsZ1 localization does not depend on any known factor. Unusually, deletion mutants of the two *ftsZ* genes were possible either individually or together, causing severe cell division defects, but the cells remained viable (6). Deletion of *ftsZ1* led to the loss of FtsZ2-ring formation, and the cells divided poorly, suggesting that FtsZ1 probably acts as a recruitment hub for other downstream cell division proteins. In the absence of FtsZ2, cells failed to divide since no constriction was observed, showing that FtsZ2 is likely involved in the later steps of cell division (6).

The membrane anchor for FtsZ in archaea was shown to be SepF, which systematically co-occurs with FtsZ in all archaea that have it (7). The archaeal SepF was shown to be essential (8), making it currently the only known FtsZ membrane anchor in archaea. In archaea with two FtsZ homologs, SepF only interacts with FtsZ2 (8). How and by what protein FtsZ1 is attached to the membrane, is currently unknown and might indicate the existence of a so far unknown additional FtsZ1 membrane anchor. Proper localization of SepF to the site of cell division depends on the presence of FtsZ1, although no direct interaction has been reported (8). In contrast to bacterial SepF, archaeal SepF dimers do not form polymers (7, 8).

In order to find additional archaeal cell division proteins, co-immunoprecipitation experiments in *H. volcanii* with HA-tagged SepF and subsequent mass spectrometric analysis showed enrichment of two proteins with a PRC-barrel domain (8). One of them has a binding site of the CdrS (small cell division regulator) transcription factor in front of its gene, which also controls expression of *ftsZ1* (11). The PRC-barrel domain is widespread and found in all domains of life. It is involved in many cellular processes - for example, it is present in the H subunit of the light-harvesting complex of purple proteobacteria and in the ribosomal maturation protein RimM (12). In both cases, the PRC-barrel domain is part of a larger protein. In contrast, the proteins with PRC-barrel domains found in Euryarchaeota are small with sizes of around 80 amino acids and only contain this domain (12). In total, three PRC-barrel domain-containing proteins were identified in *H. volcanii*, two by the SepF co-immunoprecipitation experiments, and one by a subsequent BLAST search. In this study, we showed that two of the three PRC-barrel domain-containing proteins are most likely directly involved in the FtsZ-based cell division process in *H. volcanii.* The first protein, CdpB1, localizes at the site of cell division and is essential. Exchange of the native *cdpB1* promotor with an inducible replacement allowed us to observe the cells under CdpB1-depleted conditions, resulting in a severe cell division defect. Likewise, CdpB2 also localized to the site of cell division and in contrast to CdpB1 generation of a deletion mutant was possible that generated severe division defects. In both mutant strains, localization pattern of FtsZ1, FtsZ2 and SepF were broader compared to the distinct ring-like localization at the cell centre in wild type cells. Moreover, we showed that both proteins together form a complex with SepF; however, CdpB2 does not directly interact with SepF, whereas CdpB1 does. A combination of crystal structure determination of the CdpB1/2 heterodimer and AlphaFold predictions showed B1/B2 heterodimers that chain into alternating filaments through B1:B2, B1:B1 and B2:B2 interfaces. Together with SepF binding to CdpB1, these findings suggest a role of CdpB1/2 in the membrane-proximal inclusion of the membrane anchor SepF into a polymer, one role of which it is to localize FtsZ2 filaments to the division site.

## Results

### PRC-barrel domain proteins localize to the site of cell division in *Haloferax volcanii*

We identified two PRC-barrel domain-containing proteins after immunoprecipitation experiments using the cell division protein SepF as bait (8). After BLAST searches to find more proteins with similarity within the same genome we identified a third. The genes for the three PRC-barrel domain-containing proteins (*hvo_1691, hvo_1964*, and *hvo_2019*) are present at different locations in the genome and are neither organized into operons with other genes, nor are their neighbouring genes encoding any other known cell division-related proteins (Supplementary Fig 1a). The three proteins are small, with an average size of approximately 10 kDa. They consist entirely of a PRC-barrel domain and share highly conserved residues with each other (Supplementary Fig. 1b).

To begin assessing their roles as potential new cell division proteins, we examined their localizations within cells. The three PRC-barrel domain-containing proteins identified in *Haloferax volcanii* were fused with a C-terminal mNeongreen fluorescent protein and expressed from plasmids, under the control of their native promoters. The PRC-barrel domain-containing protein from gene locus *hvo_1691* was enriched most in our pulldown experiments (8). The mNeongreen fusion protein from this locus showed strong localization at the cell center, with some additional foci at the cell periphery (Fig. 1a). It is possible that these foci are a consequence of high expression levels even though the native promotor was used. The second highest abundance of the three PRC-barrel domain proteins after immunoprecipitation with SepF is encoded by *hvo_1964.* Hvo_1964 also showed clear localization at the site of cell division, but no foci (Fig. 1c). Because of their division site localization pattern, we named these proteins <u>c</u>ell <u>d</u>ivision <u>p</u>roteins B1, CdpB1 (*hvo_1691*), and CdpB2 (*hvo_1964*). As mentioned, sequence similarity searches identified another protein with a PRC-barrel domain, hereafter called CdpB3 (*hvo_2019*), which was not found in the SepF immunoprecipitation fractions. In line with that, and in contrast to the other two CdpB proteins, CdpB3 showed only faint localization at the cell centre and was instead found throughout the cytoplasm (Supplementary Fig. 2a).

**Fig 1:**
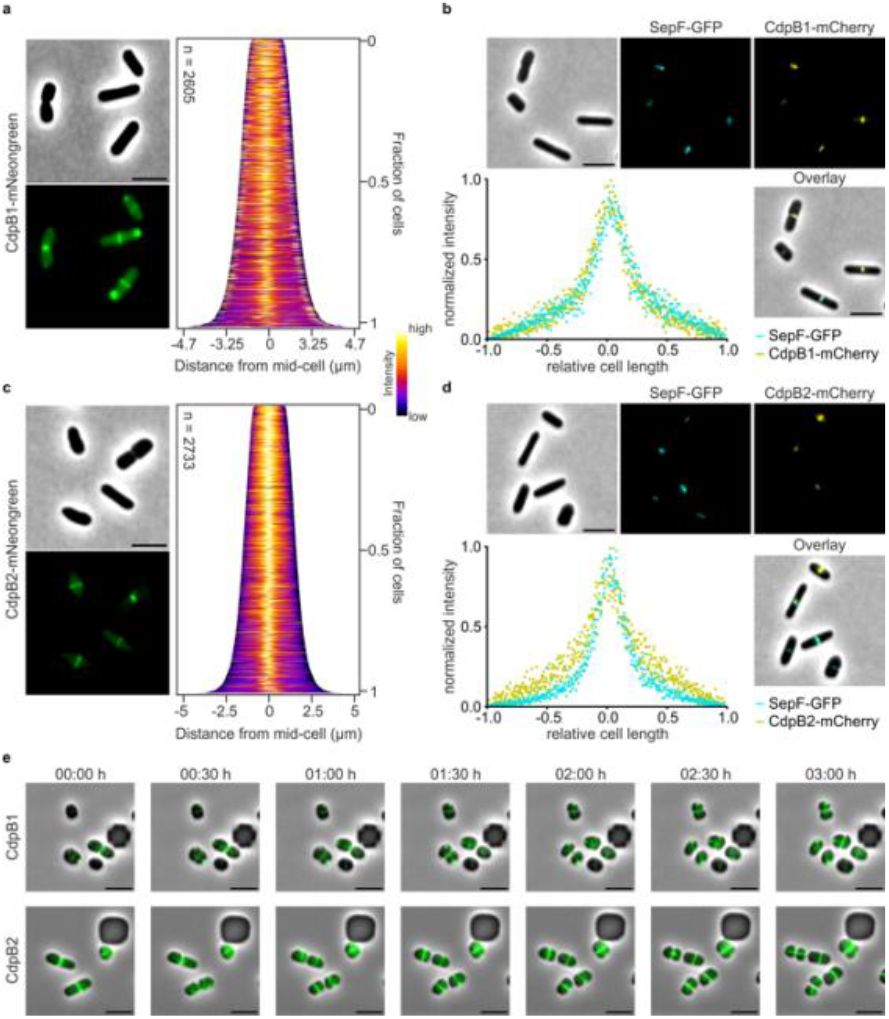
Cellular positioning of PRC barrel proteins CdpB1 and CdpB2 and their co-localization with divisome protein SepF. **a** Fluorescence microscopy of cells in early exponential phase expressing CdpB1-mNeongreen under control of the native *cdpB1* promotor. Demographic analysis shows that the highest signal intensity is at the cell centre, together with some polar foci. **b** Co-localization of SepF-GFP with CdpB1-mCherry in cells during early exponential phase. The intensity profiles of normalized SepF-GFP (cyan) and CdpB1-mCherry (yellow) show strong overlap. **c** Fluorescence microscopy of cells in early exponential phase expressing CdpB2-mNeongreen under control of the native *cdpB2* promotor. Demographic analysis shows that the proteins are localized in a ring like structure at the cell centre. **d** Co-localization of SepF-GFP together with CdpB2-mCherry in cells during early exponential phase. The intensity profiles of normalized SepF-GFP (cyan) and CdpB2-mCherry (yellow) show a broader localization for CdpB2. All localization experiments were performed in three independent replicates, with > 1000 cells used in total for analysis. Scale bar: 4 μm. **e** Time lapse microscopy of cell expressing CdpB1-/CdpB2-mNeongreen under control of their native promotors in microfluidic chambers. Movies were recorded for 16 h and a selection of 3 h of the videos are shown. For each construct at least three independent movies were recorded showing essentially the same results. Scale bar: 4 μm.

To further investigate the role of CdpB proteins during cell division in *H. volcanii*, we performed co-localization studies together with SepF. To this end, SepF-fused C-terminally with smRS-GFP was co-expressed with CdpB1- or CdpB2-mCherry from a plasmid and under the control of a tryptophan-inducible promoter. The CdpB1 signal showed strong overlap with the signal of SepF (Fig. 1b), while the CdpB2 signal appeared broader than that of SepF (Fig. 1d). However, all three proteins showed peak intensities at the cell centre. For completeness, the colocalization of SepF and CdpB3 was also investigated; however, it did not colocalize with SepF, mostly because CdpB3 was localized in the cytoplasm as reported above (Supplementary Fig. 2b). Additionally, we investigated the co-localization of the CdpB proteins with each other. CdpB1 and CdpB2 showed largely overlapping signals at the cell centre (Supplementary Fig. 2d), suggesting that they are likely involved in the cell division process at the same time and subcellular location. Again, owing to the rather diffuse distribution of CdpB3 in the cells, its signal hardly overlapped with those of CdpB1 and CdpB2 (Supplementary Fig. 2e/f).

Using live cell imaging in microfluidic plates and over complete cell cycles, CdpB1 was localized at the cell centre during the entire cycle, and immediately relocated to the site of future cell division in new-born daughter cells. The CdpB1 foci mentioned above were not involved in the cell division process but increased in intensity over time, supporting our suspicion that these might be protein aggregates rather than functional proteins. Very similar to CdpB1, CdpB2 was present at the cell centre over the complete cell cycle and immediately relocated to the future site of cell division (Fig. 1e, Supplementary Videos 1 and 2). Interestingly, CdpB3 only shortly before constriction localized to the division site and was distributed to the cytoplasm for the rest of the cell cycle (Supplementary Fig. 2c, Supplementary Video 3). Together, our results so far suggest that the CdpB1/2 proteins with a PRC domain are most likely part of the cell division machinery.

### Deletions of CdpB1 and CdpB2 have strong effects on cell shape and viability

To further illuminate the specific functions of CdpB proteins, we generated deletion strains of their genes. Attempts to delete *cdpB1* failed, indicating essentiality, possibly because of the protein’s role in cell division. To gain more insights into CdpB1’s role, we generated a conditional depletion strain by replacing its native promoter with a tryptophan-inducible promoter. This approach was used before for the essential SepF protein of *H. volcanii* (8). However, in contrast to SepF, the tryptophan promoter was too weak to allow the generation of sufficient levels of CdpB1 to restore a complete wild type phenotype of the depletion strain HTQ275 (Supplementary Fig. 3a). Yet, we were able to grow HTQ275 in full medium and then observed the effect of CdpB1 depletion upon transfer into medium lacking tryptophan. During CdpB1 depletion, the cells failed to divide efficiently and became either filamentous or enlarged with a rough surface compared to wild type cells (Fig. 2a, Supplementary Video 4 and 5). This appearance resembled cells in which FtsZ1, FtsZ2, or SepF had been deleted previously (6, 8). We therefore localized FtsZ1, FtsZ2, and SepF in the CdpB1 depletion strain HTQ275 (Fig. 2b). For quantification, we measured the areas covered by the FtsZ1, FtsZ2, and SepF signals, and compared them with the wild type area of the respective proteins. Absence of CdpB1 had an effect on the localization of all three proteins: they covered a much broader area than in the cells of our laboratory wild type strain H26, indicating that CdpB1 is required to establish a spatially constraint division site (Fig. 2c, Supplementary Table 1). Moreover, multiple division sites were visible in filamentous CdpB1 cells. In contrast to the filaments observed in the microfluidic chambers (Fig. 2a), filamentous HTQ275 cells harbouring plasmids appeared less rough (Fig 2b, Supplementary Fig. 4a). It has been observed previously that the presence of plasmids, including “empty” backbone plasmids with no additional *H. volcanii* genes to express, have a noticeable effect on the shape of cells, with impaired cell division and/or affected cell envelope growth (6, 8, 13). The reasons for this remain unclear. Finally, we used the CdpB1 depletion strain to determine whether the CdpB1-mNeongreen fusion protein is functional in cell division. Regarding cell viability, the fusion protein was able to reverse the viability defect of HTQ275 caused by CdpB1 depletion back to the wild type level. However, cell shape was not restored (Supplementary Fig. 5a-c).

**Fig 2:**
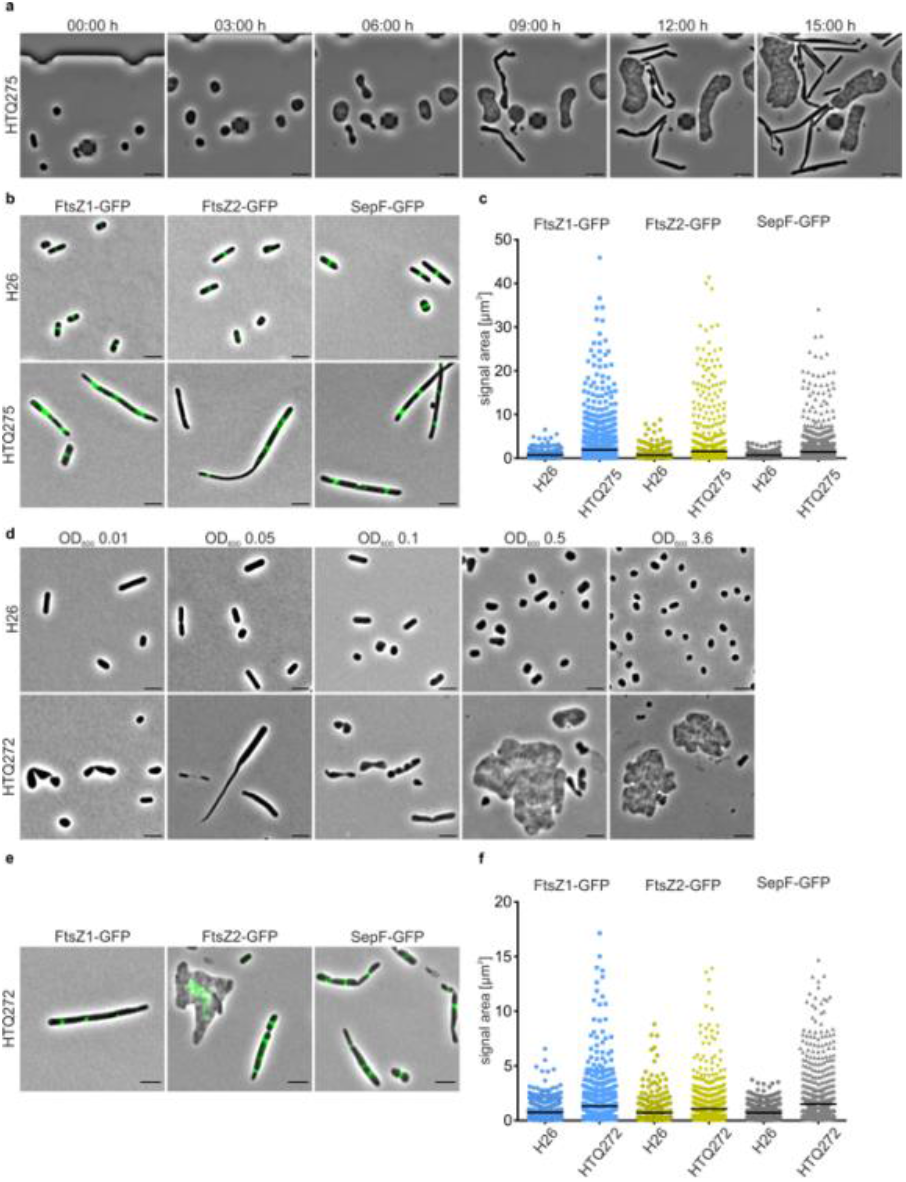
Effect of CdpB1 depletion and CdpB2 deletion on cell shape and the positioning of cell division proteins. **a** Time lapse microscopy of HTQ275 cells under CdpB1 depletion conditions in a microfluidic chamber. Only part of the 15 h movie is shown. Three independent movies were recorded showing essentially the same results. Scale bar: 4 μm. **b** Fluorescence microscopy of wild type cells (H26) and CdpB1-depleted cells (HTQ275) expressing GFP tagged, *ftsZ1, ftsZ2* and *sepF.* **c** Analysis of the signal area of each GFP construct in both strains. Each signal area per strain is plotted individually, and the mean is indicated as a black bar. Includes data from three independent replicates per strain and GFP construct, with > 1000 cells being analysed in total **d** Microscopy of the wild type (H26) and the *cdpB2* deletion strain (HTQ272) during different growth stages (optical density [OD_600_]: 0.01 lag-phase, 0.05-0.5 exponential phase, 3.6 stationary phase). **e** Fluorescence microscopy of the *cdpB2* deletion strain HTQ272 expressing GFP tagged, *ftsZ1*, *ftsZ2* and *sepF*. **f** Analysis of the signal area of each GFP construct in H26 and HTQ272. Each signal area per strain is plotted individually, and the mean is indicated as a black bar. Includes data from three independent replicates per strain and GFP construct, with > 1000 cells being analysed in total Scale bar: 4 μm.

In contrast to CdpB1, it was possible to obtain a *cdpB2* knockout strain, HTQ272, and its cells showed reduced viability compared to the wild type (Supplementary Fig. 3b). During growth, *H. volcanii* undergoes a morphological transition from rod-shaped cells during the early exponential phase to plate-shaped cells approaching the mid-exponential phase and eventually the stationary phase (14). Interestingly, the *cdpB2* deletion strain showed different cell shapes compared to the wild type, starting as filamentous cells during early growth and ending up very enlarged at the end of the exponential phase (Fig. 2d), several times larger than the wild type cells. Since the *cdpB2* deletion cells had a variety of cell shapes and were not only filamentous during the complete growth, we quantified the cell area instead of the cell length to compare them with the wild type cells (Supplementary Fig. 4b; Supplementary Table 2). We then assessed the localization of other cell division proteins of *H. volcanii* in the *cdpB2* knockout strain HTQ272 as we did for HTQ275, the *cdpB1* knockout. Similar to the CdpB1 depletion strain, the cells became filamentous due to the presence of the expression plasmids, with several sites of cell division (Fig. 2e). Although the localization of SepF, FtsZ1 or FtsZ2 at the cell centre was somewhat broader than in wild type cells, it was less so than in the CdpB1 depletion strain (Supplementary Table 1). And again, the CdpB2-mNeongreen fusion protein restored viability in HTQ272 cells, but not cell shape (Supplementary Fig. 5d-f).

Finally, we created HTQ271, a *cdpB3* deletion strain, but no phenotype was observed with regards to viability (Supplementary Fig. 3b), cell shape (Supplementary Fig. 4c, Supplementary Table 2) or localization of FtsZ1, FtsZ2, or SepF, all compared to wild type H26 (Supplementary Fig. 6a, b, Supplementary Table 1). During growth, cell shape resembled rods of the wild type during the early exponential phase and became plate-shaped as growth continued. Also, the other cell division proteins were localized in a well-defined ring at the cell centre, as observed for the wild type. The double deletion strain of both *cdpB2* and *cdpB3*, HTQ273, showed phenotypes with regards to growth, cell shape and protein localization similar to the *cdpB2* single deletion strain (Supplementary Fig. 3b, Supplementary Fig. 4d, Supplementary Table 2 and Supplementary Fig. 6c, d), together with the other data strongly suggesting that CdpB3 does not play a major role in the cell division process, despite its localization to the site of cell division shortly before septum closure. For an overview of the signal areas of the three labelled cell division proteins in the *cdpB* deletion/depletion strains they are plotted together in Supplementary Figure 6 and the mean areas are listed in Supplementary Table 2.

### Correct localization of CdpB proteins depends on the presence of other cell division proteins

Previous studies revealed that the assembly of the *H. volcanii* cell division machinery follows a seemingly hierarchical order. FtsZ1 needs to be present for the correct localization of SepF (8) and FtsZ2 (6). Additionally, SepF must be there for the assembly of the FtsZ2 ring (8). To gain further insight into the specific roles of CdpB proteins and their interplay with other cell division proteins during division, we assessed their localization in the absence of SepF, FtsZ1 or FtsZ2. SepF of *H. volcanii* was shown to be essential (8); thus, the localization of CdpB proteins was analysed in a SepF-depleted strain (8), HTQ239. Imaging was initiated immediately after the cells were transferred into medium lacking tryptophan, to induce SepF depletion. At that point, before depletion, HTQ239 had the same cell shape as the control, and CdpB1 and CdpB2 showed the same localization patterns as those observed in wild type cells (Fig 3a, b). As SepF depletion progressed, the cells became filamentous, similar to the CdpB1 depletion strain. Filamentation of the SepF-depleted strain HTQ239 in the presence of a plasmid was described previously (8) and was not the result of additional expression of CdpB1- or CdpB2-mNeongreen. After 6 h of SepF depletion, CdpB1 was hardly localized in ring-like structures, whereas CdpB2 was still sporadically localized at the site of cell division. Even 9 h after SepF depletion, CdpB2 ring-like structures were still visible; however, more foci were also found. Finally, 24 h after the initiation of SepF depletion, only CdpB1 and CdpB2 foci were observed, whereas both proteins were still localized in a ring-like structure at the cell centre in the control strain (Fig 3a, b). For CdpB3, faint localization at the cell centre was immediately lost upon the initiation of SepF depletion. However, in contrast to CdpB1 and CdpB2, free CdpB3 did not form foci but was found throughout the cytoplasm during SepF depletion (Supplementary Fig. 7a).

**Fig 3:**
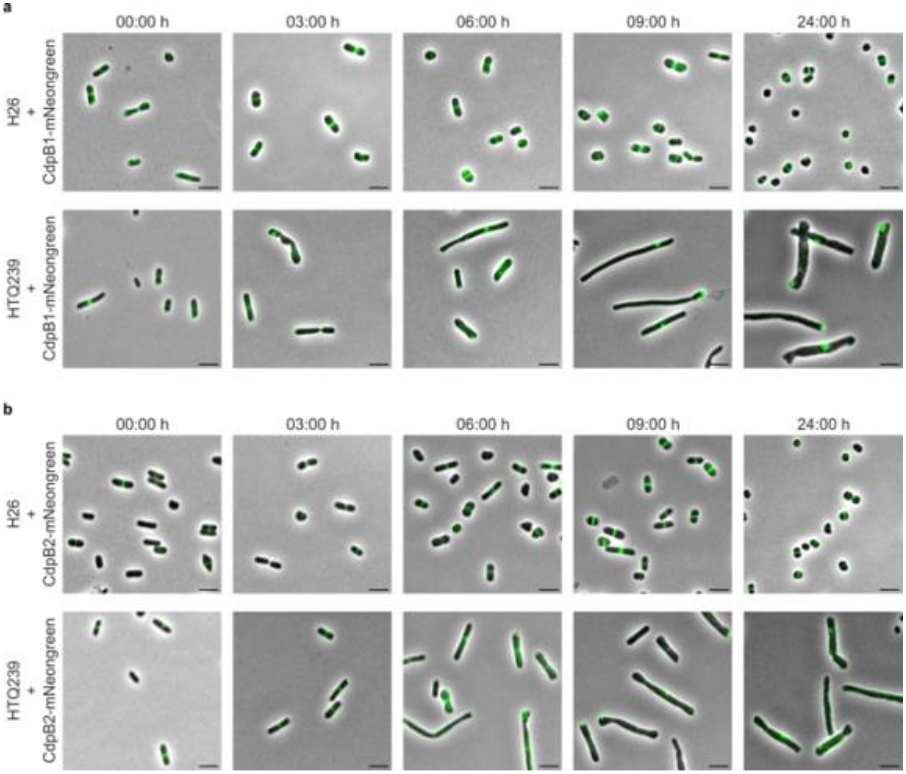
CdpB1 and CdpB2 localization during SepF depletion. **a** Fluorescence microscopy of H26 and the SepF depletion strain HTQ239 expressing CdpB1-mNeongreen from a plasmid under the control of its native promotor. Cells were imaged before (0 h) and at different timepoints after SepF depletion was induced. **b** Fluorescence microscopy of H26 and the SepF depletion strain HTQ239 expressing CdpB2-mNeongreen from a plasmid under the control of its native promotor. Cells were imaged before (0 h) and at different timepoints after SepF depletion was induced. The experiments were repeated three times with essentially the same outcome. Scale bar: 4 μm.

Although the *ftsZ1* deletion strain was transformed with a plasmid, no filamentation occurred but enlarged, rough cells appeared that made it difficult to determine the localization of the labelled CdpB proteins (Supplementary Fig. 7b). For CdpB1 and CdpB2, diffuse localization throughout the cell and additional foci scattered across the cells were observed, indicating that the absence of FtsZ1 strongly influences CdpB1 and CdpB2 positioning. CdpB3 was, as in the SepF depletion strain, observed throughout the cells. The *ftsZ2* deletion strain exhibited filamentation upon transformation with CdpB-mNeongreen expression plasmids. The defined aspect ratios of *ftsZ2* deletion cells allowed us to observe the localization of CdpB proteins accurately. Despite some foci formation of CdpB1 and CdpB2, all three CdpB proteins were localized in a ring-like structure, sometimes in more than one (Supplementary Fig. 7b). Cells of the *ftsZ1/ftsZ2* double deletion mutant were not as enlarged as those of the single *ftsZ1* deletion mutant but were also not filamentous. The CdpB proteins were not organized in a ring-like structure and only CdpB1 foci were observed, while the other two CdpB proteins were diffusely distributed in the cells (Supplementary Fig. 7b).

Moreover, to see if interdependency between CdpB1 and B2 positioning exists, we investigated the localization of CdpB2 in the CdpB1 depletion strain and vice versa the CdpB1 localization in the CdpB2 deletion strain. Indeed, proper localization of CdpB2 depends on CdpB1 but CdpB1 formed a ring like structure at the septum independently of CdpB2 (Supplementary Fig. 7d, e).

We conclude that CdpB1 localization depends on the presence of SepF, whilst CdpB2 positioning depends on the correct localization of CdpB1.

### CdpB1 and CdpB2 together form a complex with SepF

The loss of CdpB1 and CdpB2 localization during SepF depletion and the dependency of CdpB2 on CdpB1 for correct localization suggests interdependency between SepF and CdpB proteins. To study their interactions, and because *H. volcanii* is a halophilic organism, we switched to biochemically highly tractable homologs of CdpB1, CdpB2, and SepF from the hyperthermophilic archaeon, *Archaeoglobus fulgidus*. These proteins were heterologously expressed in *E. coli* with a N-terminal His-SUMO-tag, purified, the His-SUMO-tag cleaved off and analysed by size-exclusion chromatography (SEC) in different combinations. CdpB1 and CdpB2 alone showed dimer formation, as did SepF. The combination of CdpB1 and SepF led to a shift in the elution profile to smaller volumes, most likely indicating complex formation (Fig. 4a). While CdpB2 did not interact with SepF (Fig. 4b), CdpB1 and CdpB2 formed a complex (Fig. 4c). Moreover, when all three proteins were mixed, they eluted together in one peak, suggesting the formation of a tripartite SepF-CdpB1-CdpB2 complex (Fig. 4d). An SDS-PAGE of the peak elution fractions of the individual proteins and their complexes is shown in Supplementary Fig 8a. We were also able to demonstrate tripartite complex formation of SepF, CdpB1 and CdpB2 using *H. volcanii* proteins (Supplementary Fig. 8b-e).

**Fig 4:**
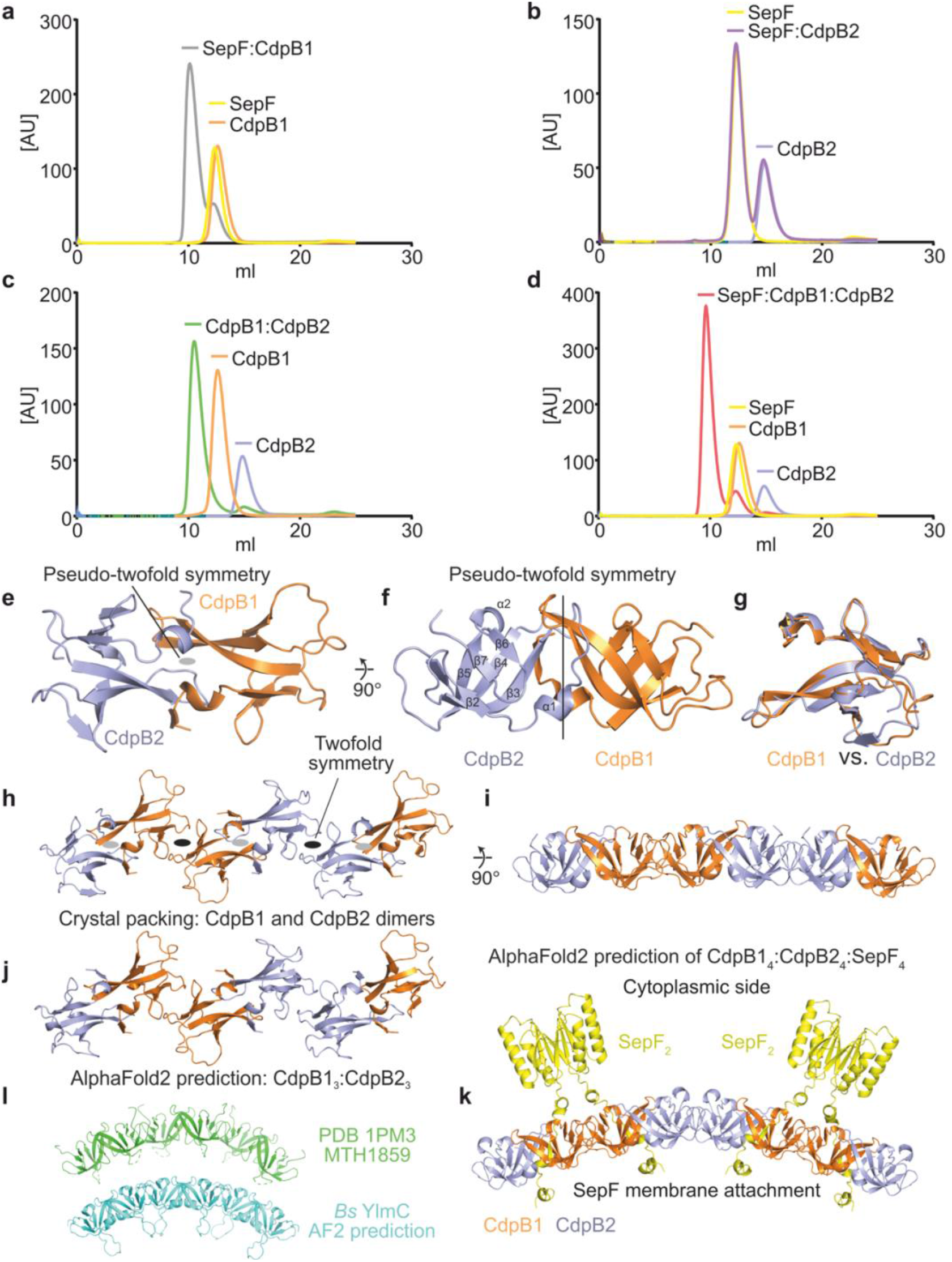
Biochemical analysis of CdpB1, CdpB2 and SepF from *Archaeoglobus fulgidus* and crystal structure of the CdpB1:CdpB2 complex. a-d Size exclusion chromatography to investigate complex formation. **a** SepF (yellow) and CdpB1 (orange) alone and SepF:CdpB1 together (grey). **b** SepF (yellow) and CdpB2 (blue) alone and SepF:CdpB2 together (purple). **c** CdpB1 (orange) and CdpB2 (blue) alone and CdpB1:CdpB2 together (green). **d** SepF (yellow), CdpB1 (orange) and CdpB2 (blue) alone and SepF:CdpB1:CdpB2 together (red). **e** Crystal structure of the CdpB1 (orange) and the CdpB2 (blue) heterodimer at 2.3 Å resolution (top view showing the pseudo-twofold symmetry axis). **f** CdpB1:CdpB2 structure side view. **g** Superposition of the CdpB1 and CdpB2 monomers showing their strong similarity. **h** Crystal packing of the CdpB1 and CdpB2 homodimers revealing alternating filaments of B1 and B2 homo- and heterodimers (top view, same as e). **i** Same as h, but side view revealing a slight bend (side view same as f). **j** AlphaFold2 predicts the same alternating filament of CdpB1:CdpB2. **k** AlphaFold2 prediction of CdpB1:CdpB2 together with SepF (yellow). SepF’s N-terminal amphipathic helix is located at the concave side of the filament, whereas the FtsZ-binding dimeric C-terminal SepF domain is on the convex side, which is presumed to face the cytoplasm of the cell where FtsZ would be located. **l** Previous crystal structure of putative adapter protein MTH1859 from *Methanobacterium thermoautotrophicum* (green, top) and AlphaFold2 prediction of YlmC from *Bacillus subtilis* (cyan, bottom). Both reveal an analogous homomeric filament to the one formed by CdpB1:CdpB2 and shown in h & i.

### Crystal structure of the CdpB1 and CdpB2 heterodimer

To obtain further insights into SepF:CdpB1:CdpB2 interactions, we attempted to crystallize the tripartite complex using *A. fulgidus* proteins. After some trials we could only obtain crystals of the CdpB1:CdpB2 complex. Crystals diffracted to 2.3 Å resolution, were orthorhombic with space group P2_1_2_1_2_1_ and contained four heterodimers of CdpB1:B2 in the asymmetric unit. The structure was solved by molecular replacement using an AlphaFold2 model of CdpB1. Crystallographic and refinement statistics are summarized in Supplementary Table 3. For two heterodimers, residues 1-92 and 1-76 were built for CdpB1 and B2, respectively. In the other two, only residues 1-76 were visible in the electron density for both CdpB1 and B2. All four heterodimers are very similar to each other and will be discussed together (Fig. 4e, f). The B1:B2 heterodimer is built through pseudo-twofold symmetry that relates the two PRC domains to each other. CdpB1 and B2 are very similar in structure, including their (small) loop regions (Fig. 4g). They adopt near-identical all-β-folds, similar to *Methanobacterium thermoautotrophicum* MTH1859 (PDB ID 1PM3) (15). Briefly, their mainly anti-parallel six-stranded β-sheet is heavily bent to form a U-structure, with the two halves of the sheet connected through strand β3 and sandwiched onto each other (Fig. 4f). The head-to-head dimerization interface is comprised mainly by strands β3 and β4 symmetrically crossing over between the two subunits. A tiny strand, β1, followed by a short one-turn α-helix, α1, extends the β-sheet of the partner subunit by packing anti-parallel against the C-terminus of β3, reinforcing the dimerization interface (Fig. 4e, f).

### Crystal packing and AlphaFold suggest CdpB1 and CdpB2 filament formation

Inspection of the crystal packing reveals that there are also homomeric interactions present, both for CdpB1:CdpB1 and for CdpB2:CdpB2, and they use two-fold symmetry in very similar ways (Fig. 4h). This arrangement of pseudo-twofold CdpB1:B2 heterodimers and two-fold CdpB1:B1 and B2:B2 homodimers creates open-ended symmetry, meaning a polymer, a filament. However, the crystals do not contain very long filaments because the filament is bent in one direction (Fig. 4i). Since crystal symmetry is used to form the filament, the filaments cannot extent all the way through the orthorhombic lattice of the crystals. AlphaFold2 predictions using CdpB1 or B2 monomers reproduce the crystal structure near-perfectly, and reveal the same heterodimer as revealed experimentally with high precision. Giving more B1 and B2 monomers to AlphaFold2 predicts the same alternating polymer as the one revealed in the crystal packing (Fig. 4h vs 4j). Given the above we think it is highly likely that CdpB1 and B2 form alternating filaments of heterodimers.

Given that AlphaFold2 easily produced results identical to our crystal structure, we then used it to predict the structure of the complex of CdpB1:B2 with SepF, which we demonstrated biochemically using purified proteins exists (Fig. 4d, Supplementary Fig. 8e). For this we used several copies of each protein since B1:B2 were shown to polymerise and SepF is known to form dimers. The resulting prediction (Fig. 4k) reveals the same CdpB1:B2 alternating filament, with SepF dimers bound to CdpB1 dimers. The N-terminal aliphatic helix of SepF that is predicted to bind to the cell membrane is located on the concave side of the B1:B2 filament, whereas SepF’s C-terminal globular domain forms the expected dimer on the convex side of the filament. Previous studies have shown that the FtsZ tail binds to the C-terminal SepF dimer (7, 16, 17) and our model would hence predict that FtsZ filaments are located on the convex side, which is facing the cytoplasm.

### PRC-barrel containing proteins are widespread in Archaea

To obtain a comprehensive overview of the distribution of the PRC barrel domain containing proteins in archaea we screened a locally-maintained database of 3,175 archaeal genome sequences for their presence using the pfam domain PF05239 that includes the CdpBs. Protein sequences containing a PRC domain are widespread in archaea, being present in all Euryarchaeota and DPANN, as well as in the Asgard and TACK groups (Figure 5, Source Data). Remarkably, the distribution of PRC domain containing proteins in archaea matches that of FtsZ and SepF, strengthening a functional link between these proteins (Figure 5). While the number of PRC domain-containing proteins ranged from one to 23 copies at the taxon level, most phyla display two copies in average, except for Methanobacteria which have an average of five copies (Figure 5, Source Data). In 93% of the cases, sequences contained only PRC domains (one or multiple). These “stand-alone” PRC proteins are found in all archaeal groups. A few PRC domain-containing proteins were associated mainly with a zinc ribbon domain in a few taxa. Intriguingly, no PRC domain-containing proteins were found in the 69 analysed genomes of Lokiarchaeia, unlike the other two groups of the Asgardarchaea (Heimdallarchaeia and Odinarchaeia), where at least one PRC protein was found (Figure 5, Source Data).

**Fig 5:**
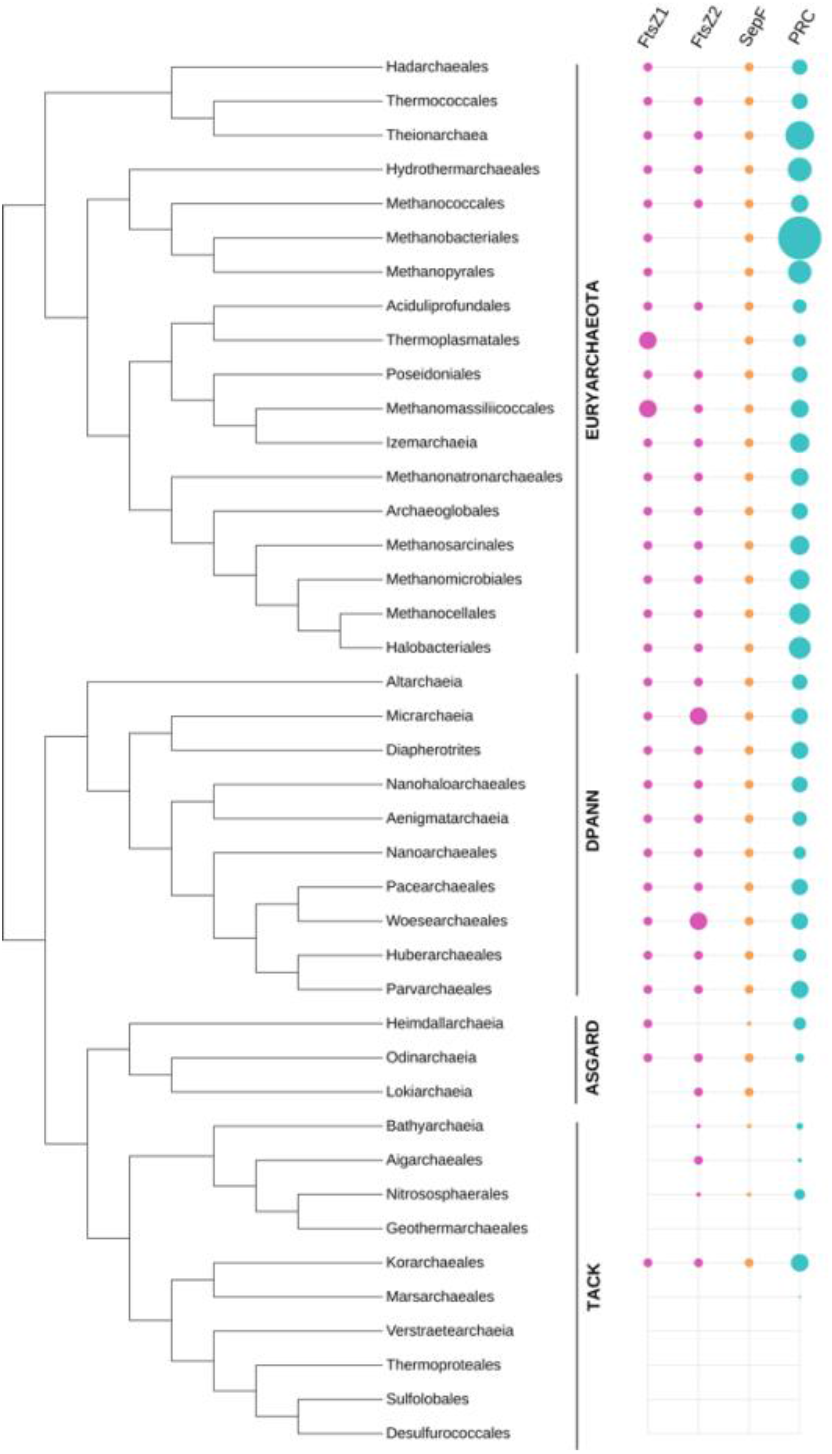
Phylogenetic distribution of PRC-domain containing proteins in archaea. Distribution of FtsZ1 (pink), FtsZ2 (pink), SepF (orange), PRC domain containing protein (turquoise) homologs on a schematic reference phylogeny of the Archaea. The schematic tree and the presence of FtsZ1, FtsZ2 and SepF are based on previous work (7). The size of the circles indicates the average number of copies found in each taxonomic group. For a complete list of these proteins, see Source Data

## Discussion

The first evidence that PRC-barrel domain-containing proteins might play a role in archaeal cell division came from an investigation of a korarchaeal genome. This *Candidatus* Korarchaeum contains a seven-gene cluster in which a PRC-barrel domain-coding gene is present, together with five *ftsZ* paralogs (18). In this study, we show that two proteins entirely consisting of this widespread PRC-barrel protein domain are actually involved in, and partially essential for FtsZ-based cell division in Archaea.

Both *Haloferax volcanii* proteins, CdpB1 and CdpB2, are important for the establishment of a distinct cell division plane in which FtsZ1, FtsZ2 and SepF are aligned in a cytokinetic ring-like structure at the cell centre. CdpB1 directly interacts with the FtsZ2 membrane anchor SepF and CdpB2, while CdpB2 only interacts with CdpB1. Together CdpB1, CdpB2 and SepF form a complex and our crystallization data suggest that CdpB1 and CdpB2 form a filament of alternating CdpB1- / CdpB2-dimers. Archaeal SepF dimers have been shown to not polymerize (7, 8) most likely due to absence of the conserved G109 that was shown to be crucial for polymerization of *Bacillus subtilis* SepF dimers (10). Bacterial SepF is associated with membrane curvature sensing and possibly membrane remodelling that requires the aforementioned polymerization of SepF dimers into oligomeric states (10, 17). The formation of filaments by the CdpB1 and CdpB2 proteins and their attachment to the membrane by SepF might support membrane activities similar to that being observed in bacteria upon SepF polymerization (Figure 6) and would make polymerization of the archaeal SepF unnecessary. Indeed, the intrinsic curvature of the CdpB1 and CdpB2 filaments resemble the one of bacterial SepF polymers at least in their direction, with negative curvature (concave side) facing the membrane (10, 17). Together with the finding that CdpB deletion and depletion experiments showed more diffuse localization of FtsZ1/2 and SepF, we suggest it is possible that CdpB1/B2/SepF polymers restrict the width of the cell constriction similar to as deduced for *B. subtilis* (10, 19) and shown in Figure 6, right for the archaeal system. Despite this intriguing model, it should be mentioned that a slight impact on vesicles by archaeal SepF dimers from *Methanobrevibacter smithii* ha*s* also been reported before (7). A next key step to verify the model would be to investigate the full complex including the FtsZ-homologs found in archaea. However, so far, we did not obtain biochemical evidence of an interaction between SepF and FtsZ, *in vitro*. Neither with the proteins of *Archaeoglobus. fulgidus*, nor with proteins from *H. volcanii*.

**Fig 6:**
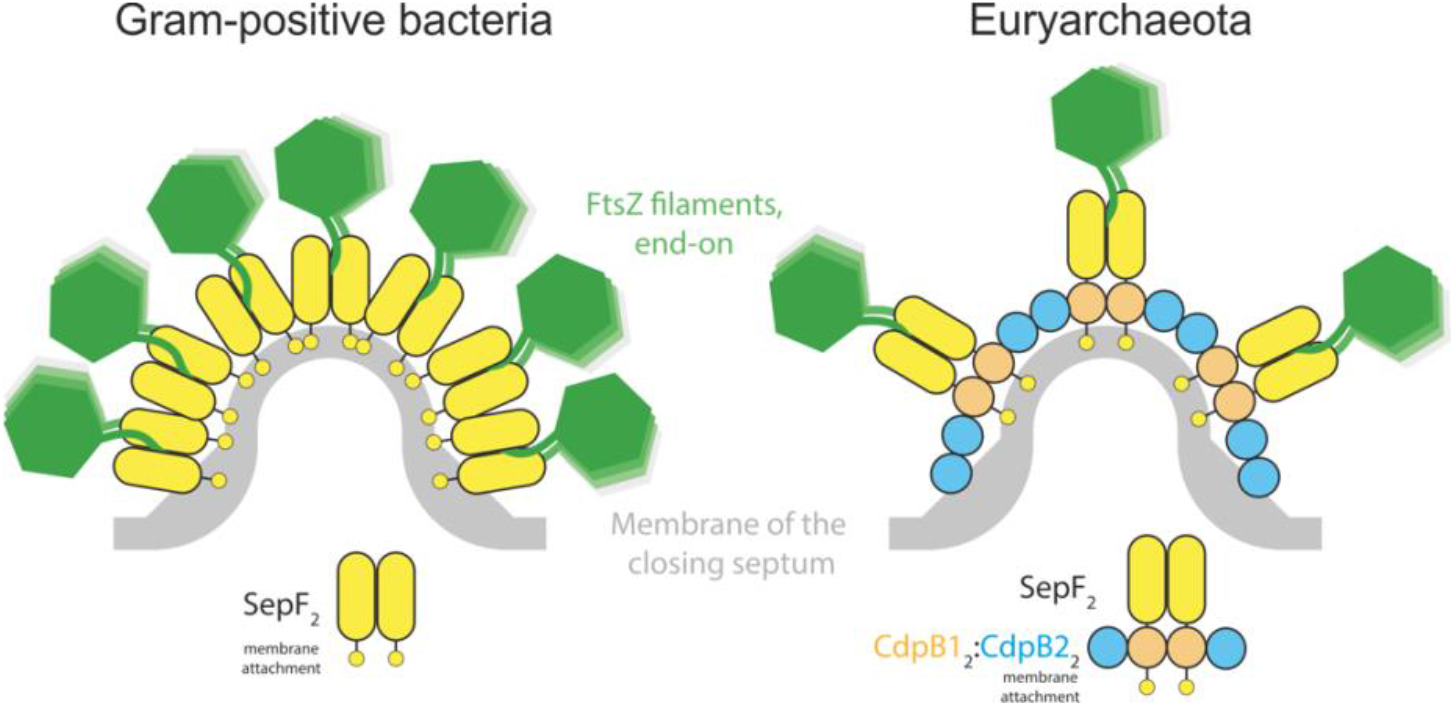
Model for CdpB1/2 function in Euryarchaeota. Left: In bacteria such as *Bacillus subtilis*, SepF dimers (yellow) polymerize into bent filaments. It has been postulated that the curvature of the SepF filaments is used to restrict the width of the nascent septum (19), which in *Bacillus* forms a cross wall. In *Bacillus*, SepF also functions as an alternative FtsZ membrane anchor, in addition to FtsA, attaching FtsZ polymers (green, running orthogonal to the 2D plane) to the cell membrane. Right: Euryarchaeal SepFs have not been shown to polymerize, which is likely due to the absence of the glycine that would be equivalent to *Bacillus* SepF G109, which is important for polymer formation. Instead, we show here that alternating units of CdpB1 (orange) and CdpB2 (blue) dimers are able to form alternating heteropolymers. We also show that SepF binds to CdpB1, which, taking the AlphaFold2 model in Fig. 4k into consideration, means that the CdpB1:CdpB2 polymer is likely attached to the cell membrane on the concave side, and to FtsZ filaments on the convex side that faces the cytoplasm. An overall arrangement analogous to the one postulated for *B. subtilis* SepF polymers emerges and it can be speculated that the B1:B2:SepF polymer might restrict cell constriction width in similar ways as in *B. subtilis.*

The assembly of the divisome of *H. volcanii* underlies a certain order and it has been shown that there is some interdependence between FtsZ1, FtsZ2 and SepF for their correct localization (6, 8). However, whether this process is hierarchically ordered as in *Escherichia coli* or rather dynamic, as in *Bacillus subtilis*, for example, is uncertain (20, 21). It has been shown that FtsZ1 has to be present at the site of cell division to ensure localization of the other two known cell division proteins to the septum (6, 8). Hence, FtsZ1 provides a platform for further downstream cell division proteins. Indeed the localization of SepF depends on the presence of FtsZ1, and in turn localization of FtsZ2 to the site of cell division relies on the presence of SepF at the septum (8). We showed that SepF also needs to be present for the formation of CdpB1 and CdpB2 in a ring-like structure at the site of cell division. Yet, mislocalization of CdpB2 might not be a direct effect of the absence of SepF, but rather of the resulting mislocalization of CdpB1, as there is no proper CdpB2 localization if CdpB1 is depleted within the cell. Interestingly, the absence of CdpB1 and CdpB2 had an impact on the complete cell division machinery, as FtsZ1, FtsZ2, and SepF were not localized into a distinct ring-like structure but, especially during CdpB1 depletion, into enlarged, unorganized foci. Moreover, it seems that also the incorporation of new S-Layer proteins, which are synthesized at the site of cell division (22) is disrupted as CdpB1 and CdpB2 mutant cells drastically increased in size in an uncontrolled manner. This increase in cell size is a general phenotype that appears upon disruption of cell division proteins in *H. volcanii* (6, 8, 23). Hence, we propose that divisome assembly in archaea that use an FtsZ based cell division is rather a dynamic process and likely not ordered.

Additionally, it seems that the PRC-barrel domain proteins are not only an important part in the divisome of archaea that divide on a FtsZ-based mechanism but also in the ESCRT-III based cell division machinery. The essential cell division protein CdvA, involved in ESCRT-III based cell division of archaea (26), contains an N-terminally located PRC-barrel domain with an so far unknown function (27, 28).

In conclusion we show that the PRC-barrel is an essential protein domain in archaeal cell division that is important for the organization of the other cell division proteins into a functional cell division machinery.

## Experimental procedures

### Material and Methods

If not stated otherwise, all chemicals were either purchased from Roth or Sigma.

### Strains and growth media

*Escherichia coli* cells for cloning and protein production were grown in lysogeny broth (LB) liquid medium or on LB-agarose plates (29) supplemented with required antibiotics. Depending on the selection marker either ampicillin (100 μg/mL) or kanamycin (25 μg/mL) were used. Cells were grown at 37 °C and liquid cultures were shaken at 150 round per minute (rpm). *Haloferax volcanii* cells were grown in full YPC-medium: 5 g/L yeast extract (Oxoid), 1 g/L Peptone (Oxoid), 1 g/L Bacto™ casamino acids (BD bioscience); pH adjusted to 7.2 with KOH (30), or selective Ca-medium: 5 g/L Bacto^TM^ casamino acids (BD bioscience); pH adjusted to 7.2 with KOH (30), supplemented with an extended trace element solution (Cab-medium) (31). If necessary, the selective medium was supplemented with 0.45 mM uracil. Liquid cultures smaller than 5 mL were grown in 15 mL tubes under constant rotation at 45 °C. Larger cultures were grown in flasks under constant shaking (120 rpm) at 45 °C. Plates to grow *H. volcanii* transformants were prepared as described before (30). To prevent evaporation plates were incubated in plastic containers at 45 °C. Strains used in this study are listed in Supplementary Table 4.

### Plasmid construction

Plasmids were either cloned via restriction enzyme-based cloning or by in vivo ligation (32). All enzymes for DNA amplification (PHUSION® polymerase) and plasmid construction (restriction enzymes, T4-DNA ligase and Quick CIP) were obtained from New England Biolabs (NEB) and used according to the manufacturer’s protocol. Plasmids for the expression of fluorescently tagged proteins were constructed via restriction enzyme-based cloning. For the knock-out plasmids around 500 base pairs (Bp) of the upstream region and around 500 Bp of the downstream region of the respective *cdpB* genes were amplified. The two fragments were connected via a *bamHI* restriction site and the ligated product cloned into linearized pTA131 (30) by in vivo ligation using 17 Bp of homologous regions per site. For the CdpB1 depletion plasmid, *cdpB1* was cloned into pTA1369 (33) to have the tryptophan-inducible promotor in front of the gene. The complete cassette of *ptnA1-cdpB1* and an *hdrB* selection marker was then cut out from the plasmid with BglII and cloned in between the up- and downstream region of the *cdpB1* knock-out plasmid, opened with BamHI. Plasmids for the heterologous expression of CdpB1, CdpB2 and SepF from *Archaeoglobus fulgidus* were constructed with in vivo ligation. The genes were ordered codon-optimized from GenScript and amplified with 15 Bp overhangs complementary, per site, to the linearized His-SUMO expression plasmid pSVA13429. Cell division proteins of *H. volcanii* for heterologous protein expression were also cloned in expression plasmid pSVA13429 as described before. All plasmids of this study, the used primers and enzymes for their construction are listed in Supplementary Table 5 and Supplementary Table 6. A table with gene sequences and the respective protein sequence of all protein constructs used in this study can be found in the Source data.

### Plasmid transformation into *H. volcanii*

The genetic system of the *H. volcanii* parent strain (H26) we were using is based on a PyrE selection marker that causes uracil auxotrophy (30). Plasmids that were transformed into *H. volcanii* were passed through a dam-/dcm- *E. coli* strain before. To transform *H. volcanii*, polyethylene glycol 600 (PEG 600) was used (30). Cells were grown to an optical density (OD_600_) of 0.8 in 10 mL and harvested (3000 x g, 10 min). The pellet was washed in 2 mL buffered spheroblast solution (1 M NaCl, 27 mM KCl, 50 mM Tris-HCl and 15 % w/v sucrose, set to pH of 8.5 after components were dissolved) and resuspended in 600 μL buffered spheroblasting solution afterwards. For one transformation, 200 μL of the prepared cell suspension was used. For spheroblast formation, 50 mM EDTA (pH 8.0) was added and the cells were incubated for 10 min at room temperature. Unmethylated plasmids (1 μg) were mixed with 83 mM EDTA and filled up to a total volume of 30 μl with unbuffered spheroblasting solution (1 M NaCl, 27 mM KCl and 15 % w/v sucrose). After 10 min, DNA was added to the cells and the tube gently inverted for mixing. Five minutes later 250 μL 60 % PEG 600 was added, the cells gently mixed and then incubated for 30 min. Next, 1.5 mL of spheroblast dilution solution (23 % salt water, 15 % w/v sucrose and 3.75 mM CaCl_2_) were added and the cells harvested at 3000 x g for 8 min. Subsequently, the cell pellet was collected at the bottom of the tubes and 1 mL of regeneration solution (18 % salt water, 1 x YPC, 15 % w/v sucrose and 3.75 mM CaCl_2_) was added to the undissolved pellet. The pellet was incubated for 1.5 h at 45 °C, dissolved by tapping the tube and then incubated for another 3.5 h. Next, cells were harvested (3000 x g, 8 min) and resuspended in 1 ml transformant dilution solution (18 % salt water, 15 % w/v sucrose and 3 mM CaCl_2_). 100 μL of transformed cells were plated on selective Ca-plates or Ca-plates supplement with 0.5 mM tryptophan if required.

### Construction of the CdpB1 depletion strain

For the generation of a tryptophan-inducible allele of *cdpB1*, integrative plasmid pSVA13510 was transformed into *H. volcanii* strain H98 (Δ*pyrE*, Δ*hdrB*). To check for correct upstream integration and correct orientation of the construct, transformants were screened via colony PCR. One colony with the correct insertion was then transferred to 5 mL YPC-medium and grown over night to induce the pop-out of the integrative plasmid. The next day the culture was diluted 1:500 into fresh YPC-medium and grown over night. This process was repeated another time to ensure pop-out of all integrated plasmids. To select for pop-out events, 100 μL of cells diluted 10^-1^,10^-2^ and 10^-3^ were plated on Ca-plates, supplemented with 50 μg/mL 5-fluoroorotic acid (5-FOA), 0.09 mM uracil, 0.5 mM tryptophan and incubated at 45 °C until colonies were visible. Then colonies were picked and transferred to a Ca-plate supplemented with uracil and 0.5 mM tryptophan. Subsequently grown colonies were transferred with a blotting paper onto a Ca-uracil plate. Four colonies that showed limited growth on the plate without tryptophan were screened via colony PCR for the correct mutation. The PCR product of one colony was sequenced and confirmed successful promoter exchange.

### Construction of the CdpB2 and CdpB3 deletion strain

For the generation of the deletion strains, either plasmid pSVA13521 (*cdpB2* deletion plasmid) or pSVA13522 (*cdpB3* deletion plasmid) were transformed into *H. volcanii* strain H26. One colony per knock-out attempt was picked and transferred into 5 mL YPC-medium to induce pop-out of the plasmid as described above. Pop-out cultures were plated in the same dilution as above on Ca-plates supplemented with 50 μg/mL 5-fluoroorotic acid (5-FOA) and 0.09 mM uracil. Possible knock-out colonies were transferred to non-selective YPC-plates and screened via colony PCR after they grew. Successful gene deletions were confirmed by sequencing. The *cdpB2*/*cdpB3* double deletion strain was constructed by transforming pSVA13523 in the *cdpB2* deletion strain, following the steps described above.

### Spot survival assay

To find the best tryptophan concentration for the CdpB1 depletion strain to grow, a spot dilution assay was performed with Ca-plates containing 0.45 mM uracil and varying tryptophan concentrations (0, 0.25, 0.5, 0.75, 1 and 1.25 mM). The preculture was grown in YPC medium overnight at 45 °C. The next day the culture was diluted to an OD_600_ of 0.2 and a serial dilution up to 10^-5^ was prepared. Of each dilution, 5 μL were spotted on the plate. The strain H26 was included as a control. Plates were sealed in a plastic bag and incubated for three days at 45 °C. Viability of the *cdpB2, cdpB3* and *cdpB2/cdpB3* deletion strains was tested with Ca-plates with no additional uracil. To overcome the uracil auxotrophy, the deletion strains and the control were transformed with empty expression vector pTA1392 complementing the *pyrE* deletion. Precultures were grown in Cab-medium and treated as the CdpB1 depletion strain the next day. Spot dilution assays to check for the functionality of CdpB1-mNeongreen and CdpB2-mNeongreen were performed as described for the deletion strains.

### Microscopy and image analysis

To investigate the cellular localizations of proteins in the wild type or mutant strains, phase contrast and fluorescence light microscopy was used. *H. volcanii* cells were transformed with respective expression plasmids and precultures of the transformants grow in 5 mL Cab-medium. The next day cells were diluted into 20 mL Cab-medium and grown until they reached an OD_600_ of 0.03. To immobilize cells, they were spotted on a 0.3 % w/v agarose pad containing 18 % salt water (144 g/L NaCl, 18 g/L MgCl_2_·6H_2_O, 21 g/L MgSO_4_·_7_H_2_O, 4.2 g/L KCl, 12 mM Tris/HCl, pH7.5). When the sample dried cells were covered with a cover slide and observed with an inverted microscope (Zeiss Axio Observer.Z1, controlled via Zeiss Blue Version 3.3.89).

To investigate the effect of the modifications and perturbations on cell shapes, precultures of the deletion strains and H26 were grown in 5 mL Cab-medium supplemented with 0.45 mM uracil at 45 °C. The next day cells were diluted into 20 mL fresh Cab-medium with uracil and grown over night again. Samples were taken at different growth stages starting at an OD_600_ of 0.01 and the last sample was taken at an OD_600_ of 3.6. Cells were imaged on agarose pads as described before.

To image the SepF depletion strain, HTQ239 transformed with different expression plasmids cells were grown in 20 mL Cab-medium, supplemented with 1 mM tryptophan, to an OD_600_ of 0.02. To induce SepF depletion, the cells were harvested (3000 x g, 10 min) and resuspended in prewarmed Cab-medium. Cells were directly imaged after resuspension as well as 3, 6, 9 and 24 h after depletion was started.

The CdpB1 depletion strain transformed with different plasmids was grown in Cab-medium, supplemented with 0.5 mM tryptophan. Since the tryptophan promotor was not strong enough to bring the viability of the CdpB1 depletion strain back to wild type levels, cells grew constantly with reduced CdpB1 levels and a transfer of the cells into medium devoid of tryptophan was not necessary to induce CdpB1 depletion. Since cells were constantly depleted of CdpB1 only samples from one timepoint were taken.

Overnight microscopy was performed using the CellASIC® ONIX2 microfluidic system and B04A-03 plates. Channels were primed with medium for 10 min under constant flow at 34.5 kPa. For cell loading, overnight cultures were diluted back to an optical density of 0.05 and flown for 15 s into the chamber with 13.8 kPa. Cells were imaged for 16 h at 45 °C under a constant flow of fresh medium at 5 kPa.

To analyse cell shapes and the areas of the signals, FIJI (Version 1.54b) (34) and the MicrobeJ (Version 5.13 l) (35) plug-in were used.

### Heterologous expression and purification of *A fulgidus* CdpB1, CdpB2 and SepF

For protein production the *E. coli* Rosetta™ strain was used. Cells were transformed either with pSVA13585 (His-SUMO-CdpB1), pSVA13586 (His-SUMO-CdpB2) or pSVA13587 (His-SUMO-SepF) and grown in LB-medium, supplemented with kanamycin overnight. The next day cells were diluted back to an OD_600_ of 0.05 in 2 L fresh LB-medium containing kanamycin and grown at 37 °C. When the cells reached an OD_600_ of 0.5 protein expression was induced by addition of 0.5 mM β-D-1-thiogalactopyranoside (IPTG). Growth was continued for 3 h at 37 °C and subsequently cells were harvested (6000 x g for 20 min) at 4 °C. The cell pellets were immediately frozen in liquid nitrogen and stored at −80°C

For purifications, cell pellets were resuspended in 20 mL Buffer A (300 mM NaCl, 10 mM imidazole, 50 mM Na_2_HPO_4_, adjusted with NaH2PO4 to pH 8.0) and lysed by passing the cells four times through a French press at 1000 psi. Subsequently, cell lysates were cleared by centrifugation for 10 min at 8000 rpm (SS-34 rotor, Sorvall). To remove *E. coli* proteins, the cell lysates were then incubated at 70 °C for 20 min under constant shaking and the precipitated proteins were removed by centrifugation at 4 °C for 10 min at 14000 g. A final centrifugation step at 83,540 x g for 45 min at 4 °C was performed, before the cleared cell lysates were loaded on a 5 mL HisTrap^TM^ HP column (Cytiva), which was equilibrated with Buffer A before. To remove unbound samples, the column was washed with Buffer B (300 mM NaCl, 20 mM imidazole, 50 mM Na_2_HPO_4_, pH 8.0) until absorbance at 280 nm was low and stable. For protein elution, Buffer C (300 mM NaCl, 250 mM imidazole, 50 mM Na_2_HPO_4_, adjusted with NaH2PO4 to pH 8.0) was applied with a constant flow of 0.5 mL/min. Samples from each purification step were loaded onto 15 % SDS-PAGE gels for analysis. Elution fractions were combined, 2 mM dithiothreitol (DTT) and 0.1 % NP-40 was added and everything incubated overnight at 4 °C with 2.5 μg/mL SUMO protease (expressed from pCDB302 and purified as described before (36)). The next day, the sample was loaded on a Superdex 75 26/600 column (Cytiva), equilibrated with Buffer D (150 mM NaCl, 25 mM Tris/HCl, pH 8.0) and run with a constant flow rate of 0.75 mL/min. Elution fractions were combined, concentrated by Amicon® ultra centrifugal filter (10 kDA cutoff, Merck Millipore) and the protein concentration determined (BCA Assay Makro Kit, Serva). Purified proteins were frozen in liquid nitrogen and stored at −80°C until used.

### Oligomeric state of CdpB1, CdpB2 and complex formation with SepF

To determine the oligomeric state of the CdpB proteins, 50 μM of each protein was loaded on a Superdex 75 10/300 column (Cytiva), equilibrated with Buffer D (150 mM NaCl, 25 mM Tris/HCl, pH 8.0). Elution was for 30 min with a constant flow rate of 0.5 mL/min. To investigate complex formation of CdpB1, CdpB2 and SepF proteins were mixed in the following combinations: CdpB1 and SepF, CdpB2 and SepF, CdpB1 and CdpB2 and CdpB1, CdpB2 and SepF (all at 50 μM each) were mixed an incubated for 10 min at room temperature, before they were applied on a Superdex 75 26/60 column (Cytiva) operated with Buffer D as described above. Elution fractions of the single- and multi-protein runs were collected, concentrated via Amicon® ultra centrifugal filters (10 kDA cutoff, Merck Millipore) and applied on 10 – 20 % gradient SDS-PAGE gels.

### Crystallization of the *A. fulgidus* CdpB1/CdpB2 complex

To obtain a stochiometric CdpB1:CdpB2 protein complex, purified proteins were mixed with a molar ratio 1:2 (B1:B2) and loaded onto a Superose 6 Increase 10/300 GL size exclusion chromatography (SEC) column (Cytiva). The column was equilibrated in 25 mM Tris/HCl, 150 mM NaCl, 5 mM MgCl_2_, 1 mM TCEP, pH 8.0. Gel filtration fractions containing the CdpB1:CdpB2 complex were pooled, concentrated using Vivaspin Turbo 5 kDa centrifugal concentrators (Sartorius). Initial crystallization hits were obtained using our in-house crystallization facility (37). Crystals were grown at 19°C by sitting-drop vapour-diffusion. Crystals were obtained in 200 nl drops composed of 100 nl of crystallization reservoir solution (5 % MPD, 0.04 M MgCl_2_, 0.05 M sodium cacodylate pH 6.0) and 100 nl of protein solution at 10 mg/mL. Crystals were harvested, cryoprotected with 30 % (v/v) glycerol in the reservoir solution and flash frozen in liquid nitrogen.

### Crystal structure determination

Diffraction data were collected at Diamond Light Source (Harwell, UK) on beamline I24 at 100 K and processed with AutoPROC (38). Anisotropic diffraction limits were applied using the online server STARANISO (39). The structure of the CdpB1:CdpB2 was solved by molecular replacement with Phaser (40), using an AlphaFold2 prediction of CdpB1 monomer as the template (41, 42). Interactive model building was performed with Coot (43), refinement with REFMAC5 (44) and phenix.refine (45) and validation with Molprobity (46). Crystallographic data and model statistics are summarised in Supplementary Table 3. Figures of atomic models were prepared with PyMOL (47).

### Heterologous expression and purification of *H volcanii* CdpB1, CdpB2 and SepF

All *H. volcanii* proteins (SepF, CdpB1 and CdpB2) were constructed with His-SUMO N-terminal fusion and expressed in *Escherichia coli* BL21 (DE3) for 3 hours at 37 °C in LB medium supplemented with 100 μg/mL ampicillin, in the presence of 1 mM IPTG after reaching the OD_600_ 0.6-0.8. Cells were pelleted at 6000 x g for 45 min, resuspended in the supernatant, again centrifuged at 4000 x g for 30 min and finally flash frozen using liquid nitrogen and stored at −80 °C. Cells were thawed and resuspended in Lysis buffer (100 mM HEPES pH 7.4, 150 mM KCl, 20 mM imidazole and 1 mM DTT) supplemented with EDTA-free protease inhibitor tablets (1 per 50 ml buffer) and 1 mg/mL DNase I. The lysis was performed using Q700 sonicator equipped with a probe of 12.7 mm diameter immersed into the resuspended pellet. The sonication was done for 10 min (amplitude 40, on time 1 s, off time 4 s). Next, the cell debris was removed by centrifugation at 31000 x g for 45 min and the cleared lysate was incubated with HisPur^TM^ Ni-NTA resin (Thermo Fisher Scientific) for 1 h. Subsequently, the resin was washed with Lysis buffer (30 column volumes) and His-SUMO tagged protein eluted with an imidazole gradient elution buffers (100 mM HEPES pH 7.4, 150 mM KCl, 100 – 400 mM imidazole, 1 mM DTT). Fractions were further analysed by SDS-PAGE, pooled and dialyzed overnight against the storage buffer (100 mM HEPES pH 7.4, 150 mM KCl, 10 % glycerol, 1 mM DTT) in the presence of His-Ulp1 protease (1:100 molar ratio) for His-SUMO cleavage. In order to remove the cleaved His-SUMO tag and His-Ulp1, reverse affinity chromatography was performed using Protino Ni-IDA resin (LACTAN Macherey Nagel). After 30 min of incubation with the resin, flow through was collected and analysed by SDS-PAGE. At this point, SepF and CdpB2 were aliquoted, flash frozen in liquid nitrogen and stored at –80 °C. CdpB1 was further loaded onto a HiLoad® 16/600 Superdex 75 size exclusion column pre-equilibrated with storage buffer. Fractions were pooled, flash frozen in liquid nitrogen and stored at −80 °C. All steps were done at 4 °C and concentrations of all proteins were determined by Bio-Rad Protein Assay Dye Reagent Concentrate.

### Size Exclusion Chromatography coupled with Multi Angle Light Scattering (SEC-MALS)

Individual proteins as well as protein complexes were resolved on a Superdex 200 Increase 10/300 (Cytiva) with a flow rate of 0.5 mL/min in high salt buffer (1 M KCl, 10 mM MgCl_2_, 100 mM HEPES, pH 7.4) coupled with miniDAWN light scattering device (Wyatt) at room temperature. The peak areas were defined based on the changes in refractive index, which was used to determine the molecular weights. The analysis was done using ASTRA software (Wyatt). Individual proteins SepF, CdpB1 and CdpB2 were run at 0.9, 0.7, 0.5 mg/mL, respectively. When two proteins were pre-mixed, the concentration of each was 0.25 mg/mL while all three proteins were pre-mixed with a concentration of 17 mg/mL per protein.

### Homology searches and sequences analysis

For PRC domain containing proteins homology searches, we assembled a local databank of 3,175 archaeal genomes representatives of all major phyla available in public databases as of January 2022. We carried out an HMM-based homology search using the pfam domain PF05239, the HMMER3.3.2 package (48) and an e-value threshold of 1. Several rounds of curation were then performed to discard false positives, using additional information such as domains organization, alignments, and phylogeny. Taxonomic distribution was then mapped onto an archaeal reference phylogeny using IToL (49). The schematic reference tree of Archaea and the distribution of FtsZ1, FtsZ2, and SepF are based on (7).

## Acknowledgements

PN and SVA were supported by a Momentum grant by the VW foundation (grant number 94933). DKC and DB were supported by the Volkswagen Stiftung “Life?” programme (to JL, grant number Az 96727) and by the Medical Research Council, as part of United Kingdom Research and Innovation (UKRI), MRC file reference number: U105184326 (to JL). NT and SG wish to acknowledge support by the French Government’s Investissement d’Avenir program, Laboratoire d’Excellence “Integrative Biology of Emerging Infectious Diseases” (grant n°ANR-10-LABX-62-IBEID), and the computational and storage services (Maestro cluster) provided by the IT department at Institut Pasteur, Paris. MK and ML were supported by the Austrian Science Fund (FWF) Stand-Alone P34607.

We thank Dr. Xing Ye for providing us the HIS-SUMO expression plasmid pSVA13429. pCDB302 was a gift from Christopher Bahl (Addgene plasmid #113673; http://n2t.net/addgene:113673;RRID:Addgene_113673).

## Author contributions

PN and SVA conceived the project and performed all experiments not mentioned otherwise. CVD purified the proteins from *A. fulgidus* and performed SEC experiments. DKC and DB solved the crystal structure. MK expressed and purified proteins from *H. volcanii* and performed SEC-MALS experiments with them. NT performed phylogenetic analysis. PN and JL prepared figures and PN wrote the manuscript draft. SG, ML, JL, SVA reviewed drafts of the manuscript, supervised work and acquired funding.

## References

1. Mahone,C.R. and Goley,E.D. (2020) Bacterial cell division at a glance. J. Cell Sci., 133, 1–7.

2. Andrade,V. and Echard,A. (2022) Mechanics and regulation of cytokinetic abscission. Front. Cell Dev. Biol., 10, 1–16.

3. Makarova,K.S., Yutin,N., Bell,S.D. and Koonin,E. V. (2010) Evolution of diverse cell division and vesicle formation systems in Archaea. Nat. Rev. Microbiol., 8, 731–741.

4. Ithurbide,S., Gribaldo,S., Albers,S.-V. and Pende,N. (2022) Spotlight on FtsZ-based cell division in Archaea. Trends Microbiol., 30, 665–678.

5. Makarova,K.S. and Koonin,E. V. (2010) Two new families of the FtsZ-tubulin protein superfamily implicated in membrane remodeling in diverse bacteria and archaea. Biol. Direct, 5, 1–9.

6. Liao,Y., Ithurbide,S., Evenhuis,C., Löwe,J. and Duggin,I.G. (2021) Cell division in the archaeon Haloferax volcanii relies on two FtsZ proteins with distinct functions in division ring assembly and constriction. Nat. Microbiol., 6, 594–605.

7. Pende,N., Sogues,A., Megrian,D., Palabikyan,H., Sartori-Rupp,A., Graña,M., Rittmann,S.K.-M.R., Wehenkel,A.M., Alzari,P. and Gribaldo,S. (2021) SepF is the FtsZ-anchor in archaea, with features of an ancestral cell division system. Nat. Commun., 10.1038/s41467-021-23099-8.

8. Nußbaum,P., Gerstner,M., Dingethal,M., Erb,C. and Albers,S.V. (2021) The archaeal protein SepF is essential for cell division in Haloferax volcanii. Nat. Commun., 12, 1–15.

9. Hamoen,L.W., Meile,J.C., De Jong,W., Noirot,P. and Errington,J. (2006) SepF, a novel FtsZ-interacting protein required for a late step in cell division. Mol. Microbiol., 59, 989–999.

10. Duman,R., Ishikawa,S., Celik,I., Strahl,H., Ogasawara,N., Troc,P., Lowe,J. and Hamoen,L.W. (2013) Structural and genetic analyses reveal the protein SepF as a new membrane anchor for the Z ring. Proc. Natl. Acad. Sci., 110, E4601–E4610.

11. Liao,Y., Vogel,V., Hauber,S., Bartel,J., Alkhnbashi,O.S., Maaß,S., Schwarz,T.S., Backofen,R., Becher,D., Duggin,I.G., et al. (2021) CdrS Is a Global Transcriptional Regulator Influencing Cell Division in Haloferax volcanii. MBio, 12, e01416–21.

12. Anantharaman,V. and Aravind,L. (2002) The PRC-barrel: a widespread, conserved domain shared by photosynthetic reaction center subunits and proteins of RNA metabolism. Genome Biol., 3, RESEARCH0061.

13. Abdul Halim,M.F., Karch,K.R., Zhou,Y., Haft,D.H., Garcia,B.A. and Pohlschroder,M. (2016) Permuting the PGF Signature Motif Blocks both Archaeosortase-Dependent C-Terminal Cleavage and Prenyl Lipid Attachment for the Haloferax volcanii S-Layer Glycoprotein. J. Bacteriol., 198, 808–815.

14. Li,Z., Kinosita,Y., Rodriguez-Franco,M., Nußbaum,P., Braun,F., Delpech,F., Quax,T.E.F. and Albers,S.-V. (2019) Positioning of the Motility Machinery in Halophilic Archaea. MBio, 10, 1–17.

15. Ye,H., Chen,T.-C., Xu,X., Pennycooke,M., Wu,H. and Steegborn,C. (2004) Crystal structure of the putative adapter protein MTH1859. J. Struct. Biol., 148, 251–256.

16. Król,E., van Kessel,S.P., van Bezouwen,L.S., Kumar,N., Boekema,E.J. and Scheffers,D.J. (2012) Bacillus subtilis SepF binds to the C-terminus of FtsZ. PLoS One, 7.

17. Sogues,A., Martinez,M., Gaday,Q., Ben-Assaya,M., Graña,M., Voegele,A., VanNieuwenhze,M., England,P., Haouz,A., Chenal,A., et al. (2020) Essential dynamic interdependence of FtsZ and SepF for Z-ring and septum formation in Corynebacterium glutamicum. Nat. Commun., 11.

18. Elkins,J.G., Podar,M., Graham,D.E., Makarova,K.S., Wolf,Y., Randau,L., Hedlund,B.P., Brochier-Armanet,C., Kunin,V., Anderson,I., et al. (2008) A korarchaeal genome reveals insights into the evolution of the Archaea. Proc. Natl. Acad. Sci., 105, 8102–8107.

19. Wenzel,M., Celik Gulsoy,I.N., Gao,Y., Teng,Z., Willemse,J., Middelkamp,M., van Rosmalen,M.G.M., Larsen,P.W.B., van der Wel,N.N., Wuite,G.J.L., et al. (2021) Control of septum thickness by the curvature of sepf polymers. Proc. Natl. Acad. Sci. U. S. A., 118, 1–9.

20. Du,S. and Lutkenhaus,J. (2017) Assembly and activation of the Escherichia coli divisome. Mol. Microbiol., 105, 177–187.

21. Errington,J. and Wu,L.J. (2017) Cell Cycle Machinery in Bacillus subtilis. In Löwe,J., Amos,L.A. (eds). Springer International Publishing, Cham, pp. 67–101.

22. Abdul-Halim,M.F., Schulze,S., DiLucido,A., Pfeiffer,F., Filho,A.W.B. and Pohlschroder,M. (2020) Lipid Anchoring of Archaeosortase Substrates and Mid-Cell Growth in Haloarchaea. MBio, 11, 863746.

23. Duggin,I.G., Aylett,C.H.S., Walsh,J.C., Michie,K.A., Wang,Q., Turnbull,L., Dawson,E.M., Harry,E.J., Whitchurch,C.B., Amos,L.A., et al. (2015) CetZ tubulin-like proteins control archaeal cell shape. Nature, 519, 362–365.

24. Caldas,P., López-Pelegrín,M., Pearce,D.J.G., Budanur,N.B., Brugués,J. and Loose,M. (2019) Cooperative ordering of treadmilling filaments in cytoskeletal networks of FtsZ and its crosslinker ZapA. Nat. Commun., 10, 5744.

25. Cheng,Y.S., Brantner,C.A., Tsapin,A. and Collins,M.L.P. (2000) Role of the H Protein in Assembly of the Photochemical Reaction Center and Intracytoplasmic Membrane in Rhodospirillum rubrum. J. Bacteriol., 182, 1200–1207.

26. Blanch Jover,A. and Dekker,C. (2023) The archaeal Cdv cell division system. Trends Microbiol., 10.1016/j.tim.2022.12.006.

27. Moriscot,C., Gribaldo,S., Jault,J.-M., Krupovic,M., Arnaud,J., Jamin,M., Schoehn,G., Forterre,P., Weissenhorn,W. and Renesto,P. (2011) Crenarchaeal CdvA Forms Double-Helical Filaments Containing DNA and Interacts with ESCRT-III-Like CdvB. PLoS One, 6, e21921.

28. Samson,R.Y., Obita,T., Hodgson,B., Shaw,M.K., Chong,P.L.G., Williams,R.L. and Bell,S.D. (2011) Molecular and Structural Basis of ESCRT-III Recruitment to Membranes during Archaeal Cell Division. Mol. Cell, 41, 186–196.

29. Bertani,G. (1951) Studies on lysogenesis. I. The mode of phage liberation by lysogenic Escherichia coli. J. Bacteriol., 62, 293–300.

30. Allers,T., Ngo,H.-P., Mevarech,M. and Lloyd,R.G. (2004) Development of Additional Selectable Markers for the Halophilic Archaeon Haloferax volcanii Based on the leuB and trpA Genes. Appl. Environ. Microbiol., 70, 943–953.

31. de Silva,R.T., Abdul-Halim,M.F., Pittrich,D.A., Brown,H.J., Pohlschroder,M. and Duggin,I.G. (2021) Improved growth and morphological plasticity of Haloferax volcanii. Microbiology, 167.

32. Watson,J.F. and García-Nafría,J. (2019) In vivo DNA assembly using common laboratory bacteria: A re-emerging tool to simplify molecular cloning. J. Biol. Chem., 294, 15271–15281.

33. Braun,F., Thomalla,L., van der Does,C., Quax,T.E.F., Allers,T., Kaever,V. and Albers,S.-V. (2019) Cyclic nucleotides in archaea: Cyclic di-AMP in the archaeon Haloferax volcanii and its putative role. Microbiologyopen, 8, 1–23.

34. Schindelin,J., Arganda-Carreras,I., Frise,E., Kaynig,V., Longair,M., Pietzsch,T., Preibisch,S., Rueden,C., Saalfeld,S., Schmid,B., et al. (2012) Fiji: An open-source platform for biological-image analysis. Nat. Methods, 9, 676–682.

35. Ducret,A., Quardokus,E.M. and Brun,Y. V (2016) MicrobeJ, a tool for high throughput bacterial cell detection and quantitative analysis. Nat. Microbiol., 1, 16077.

36. Lau,Y.-T.K., Baytshtok,V., Howard,T.A., Fiala,B.M., Johnson,J.M., Carter,L.P., Baker,D., Lima,C.D. and Bahl,C.D. (2018) Discovery and engineering of enhanced SUMO protease enzymes. J. Biol. Chem., 293, 13224–13233.

37. Gorrec,F. and Löwe J. (2018) Automated Protocols for Macromolecular Crystallization at the MRC Laboratory of Molecular Biology. J. Vis. Exp., 10.3791/55790.

38. Vonrhein,C., Flensburg,C., Keller,P., Sharff,A., Smart,O., Paciorek,W., Womack,T. and Bricogne,G. (2011) Data processing and analysis with the autoPROC toolbox. Acta Crystallogr. Sect. D Biol. Crystallogr., 67, 293–302.

39. Vonrhein,C., Tickle,I.J., Flensburg,C., Keller,P., Paciorek,W., Sharff,A. and Bricogne,G. (2018) Advances in automated data analysis and processing within autoPROC, combined with improved characterisation, mitigation and visualisation of the anisotropy of diffraction limits using STARANISO. Acta Crystallogr. Sect. A Found. Adv., 74, a360–a360.

40. McCoy,A.J., Grosse-Kunstleve,R.W., Storoni,L.C. and Read,R.J. (2005) Likelihood-enhanced fast translation functions. Acta Crystallogr. Sect. D Biol. Crystallogr., 61, 458–464.

41. Jumper,J., Evans,R., Pritzel,A., Green,T., Figurnov,M., Ronneberger,O., Tunyasuvunakool,K., Bates,R., Žídek,A., Potapenko,A., et al. (2021) Highly accurate protein structure prediction with AlphaFold. Nature, 596, 583–589.

42. Varadi,M., Anyango,S., Deshpande,M., Nair,S., Natassia,C., Yordanova,G., Yuan,D., Stroe,O., Wood,G., Laydon,A., et al. (2022) AlphaFold Protein Structure Database: massively expanding the structural coverage of protein-sequence space with high-accuracy models. Nucleic Acids Res., 50, D439–D444.

43. Emsley,P., Lohkamp,B., Scott,W.G. and Cowtan,K. (2010) Features and development of Coot. Acta Crystallogr. Sect. D Biol. Crystallogr., 66, 486–501.

44. Murshudov,G.N., Vagin,A.A. and Dodson,E.J. (1997) Refinement of Macromolecular Structures by the Maximum-Likelihood Method. Acta Crystallogr. Sect. D Biol. Crystallogr., 53, 240–255.

45. Liebschner,D., Afonine,P. V, Baker,M.L., Bunkóczi,G., Chen,V.B., Croll,T.I., Hintze,B., Hung,L.-W., Jain,S., McCoy,A.J., et al. (2019) Macromolecular structure determination using X-rays, neutrons and electrons: recent developments in Phenix. Acta Crystallogr. Sect. D Struct. Biol., 75, 861–877.

46. Chen,V.B., Arendall,W.B., Headd,J.J., Keedy,D.A., Immormino,R.M., Kapral,G.J., Murray,L.W., Richardson,J.S. and Richardson,D.C. (2010) MolProbity: all-atom structure validation for macromolecular crystallography. Acta Crystallogr. Sect. D Biol. Crystallogr., 66, 12–21.

47. Schrödinger, L. & DeLano,W. (2020) The PyMOL Molecular Graphics System.

48. Johnson,L.S., Eddy,S.R. and Portugaly,E. (2010) Hidden Markov model speed heuristic and iterative HMM search procedure. BMC Bioinformatics, 11, 431.

49. Letunic,I. and Bork,P. (2019) Interactive Tree Of Life (iTOL) v4: recent updates and new developments. Nucleic Acids Res., 47, W256–W259.

