## Supplemental material for "PRC domain-containing proteins modulate FtsZ-based archaeal cell division"

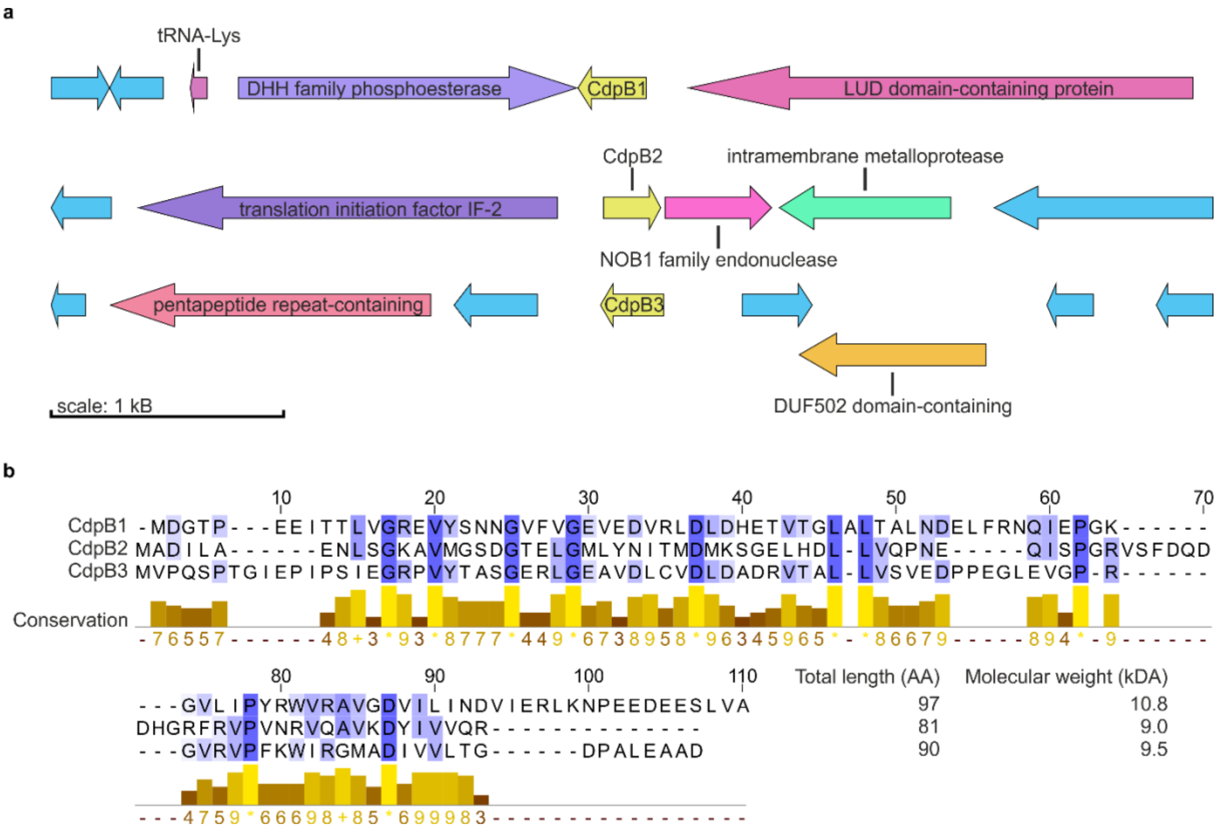


**Supplementary Figure 1: Genetic neighborhood and sequence alignment of CdpB1-3. a** Genetic neighborhood map of the *cdpB* genes: *cdpB1* (*hvo_1691*), *cdpB2* (*hvo_1964*) and *cdpB3* (*hvo_2019*). Mapped are 2500 Bp up- and downstream of the *cdpB* genes. The encoded proteins in the vicinity of the *cdpB* genes are indicated, genes that encode hypothetical proteins are shown in blue. The map was drawn with Gene Graphics (1). **b** Sequence alignment of the three CdpB proteins using the MUSCLE (Multiple Sequence comparison by Log-Expectation) (2) web service with default settings. Conserved residues are highlighted with a blue gradient additionally the conservation scores are given below the alignment using Jalview (Version 2.11.0 (3)). The total length and the predicted molecular mass for each CdpB protein are indicated as well.

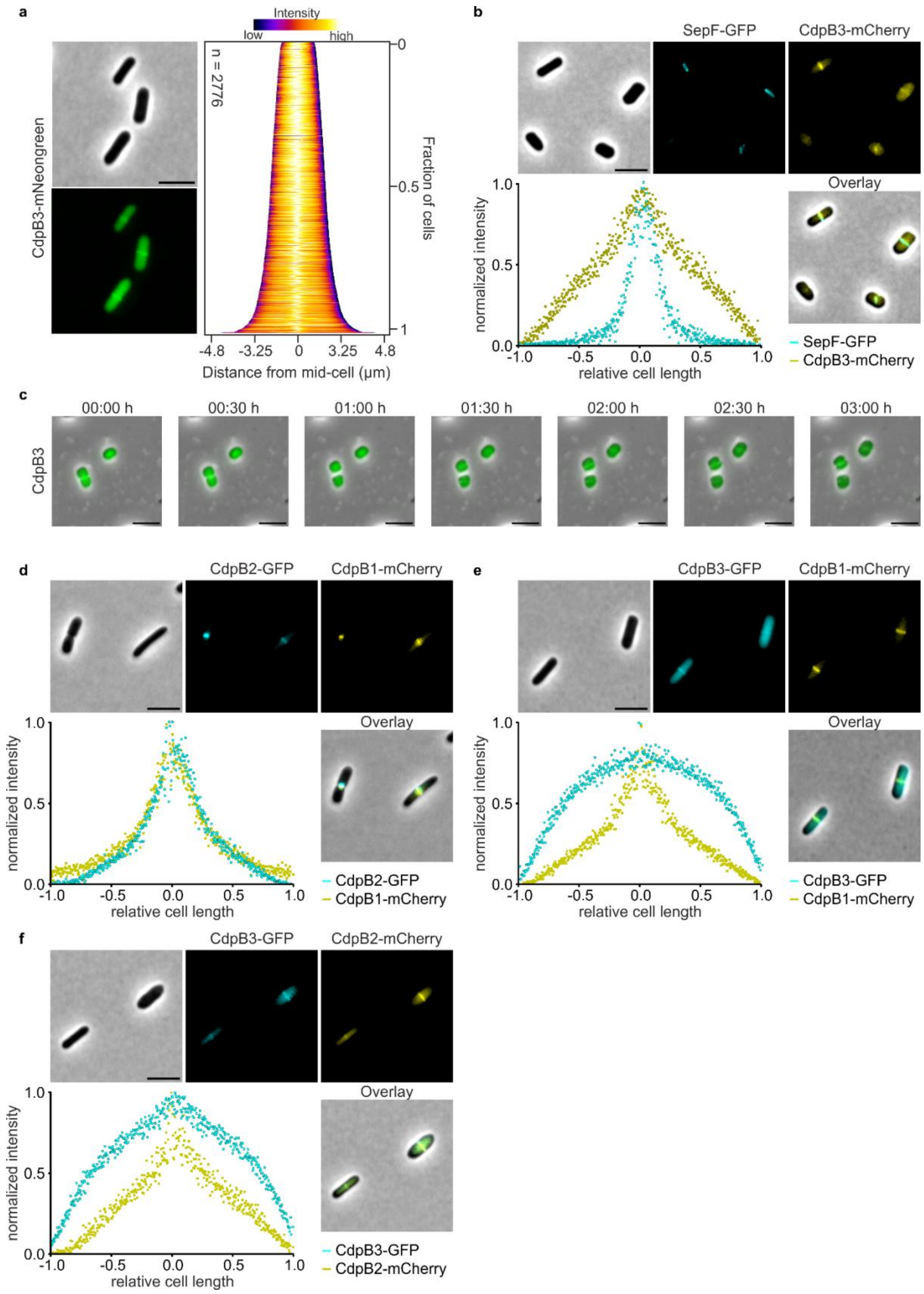


**Supplementary Figure 2: Cellular localization of CdpB3, co-localization with divisome protein SepF and co-localization of the CdpB proteins. a** Fluorescence microscopy of cells in early exponential phase expressing CdpB3-mNeongreen under control of the native *cdpB3* promotor. Demographic analysis shows that the protein is localized throughout the cytoplasm with the highest signal intensity found as a ring like structure at the cell center. **b** Co-localization of SepF-GFP together with CdpB3-mCherry in cells during early exponential phase. The intensity profile of the normalized SepF-GFP (cyan) and the CdpB3-mCherry (yellow) signal only show an overlap at the cell center. The localization experiments were performed in three independent replicates with > 1000 cells used for analysis. Scale bar: 4 µm. **c** Time lapse microscopy of cell expressing CdpB3 -mNeongreen under control of their native promotors in microfluidic chambers. In total 16 h a movie was recorded and a selection of 3 h of the video is shown. At least three independent movies were recorded showing the same results. Scale bar: 4 µm. **d** Co-localization of CdpB2-GFP together with CdpB1-mCherry in cells during early exponential phase. The intensity profile of the normalized CdpB2-GFP (cyan) and the CdpB1-mCherry (yellow) signal shows co-localization of both proteins at the site of cell division. **e** Co-localization of CdpB3-GFP together with CdpB1-mCherry in cells during early exponential phase. The intensity profile of the normalized CdpB3-GFP (cyan) and the CdpB1-mCherry (yellow) signal shows little co-localization of both proteins. **f** Co-localization of CdpB3-GFP together with CdpB2-mCherry in cells during early exponential phase. The intensity profile of the normalized CdpB3-GFP (cyan) and the CdpB2-mCherry (yellow) signal as well show little co-localization. All localization experiments were performed in three independent replicates with > 1000 cells used for analysis. Scale bar: 4 µm.

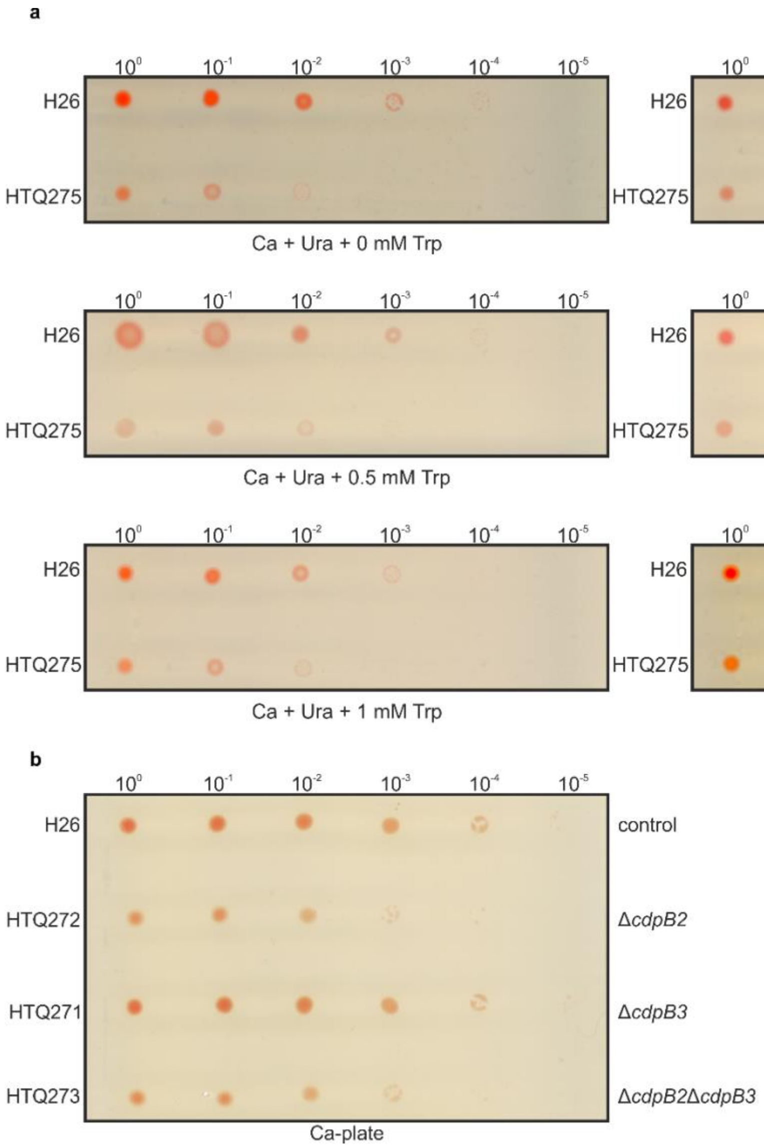


**Supplementary Figure 3: Spot dilution assay of the CdpB1 depletion strain on plates with different tryptophan concentrations and spot dilution assay of the Δ*cdpB2*, Δ*cdpB3* and Δ*cdpB2*Δ*cdpB3* strains. a** Based on a starting $OD_{600}$ of 0.2 a serial dilution of H26 and the CdpB1 depletion strain HTQ275 until $10^{-5}$ was made. The diluted cells were spotted on Ca-plates supplemented with uracil and different tryptophan concentrations to find the tryptophan concentration that complements the viability defect of CdpB1 depletion. However, the *tnaA1* promotor is not strong enough to fully complement the HTQ275 viability back to wildtype levels. Three independent replicates of the experiments were made with same results. **b** Based on a starting $OD_{600}$ of 0.2 a serial dilution of strain H26, HTQ272 (Δ*cdpB2*), HTQ271 (Δ*cdpB3*) and HTQ273 (Δ*cdpB2*Δ*cdpB3*) until $10^{-5}$ was made. The diluted cells were spotted on Ca-plates. Strain HTQ272 and HTQ273 show reduced viability compared to the wildtype H26. Three independent replicates of the experiments were made with same results.

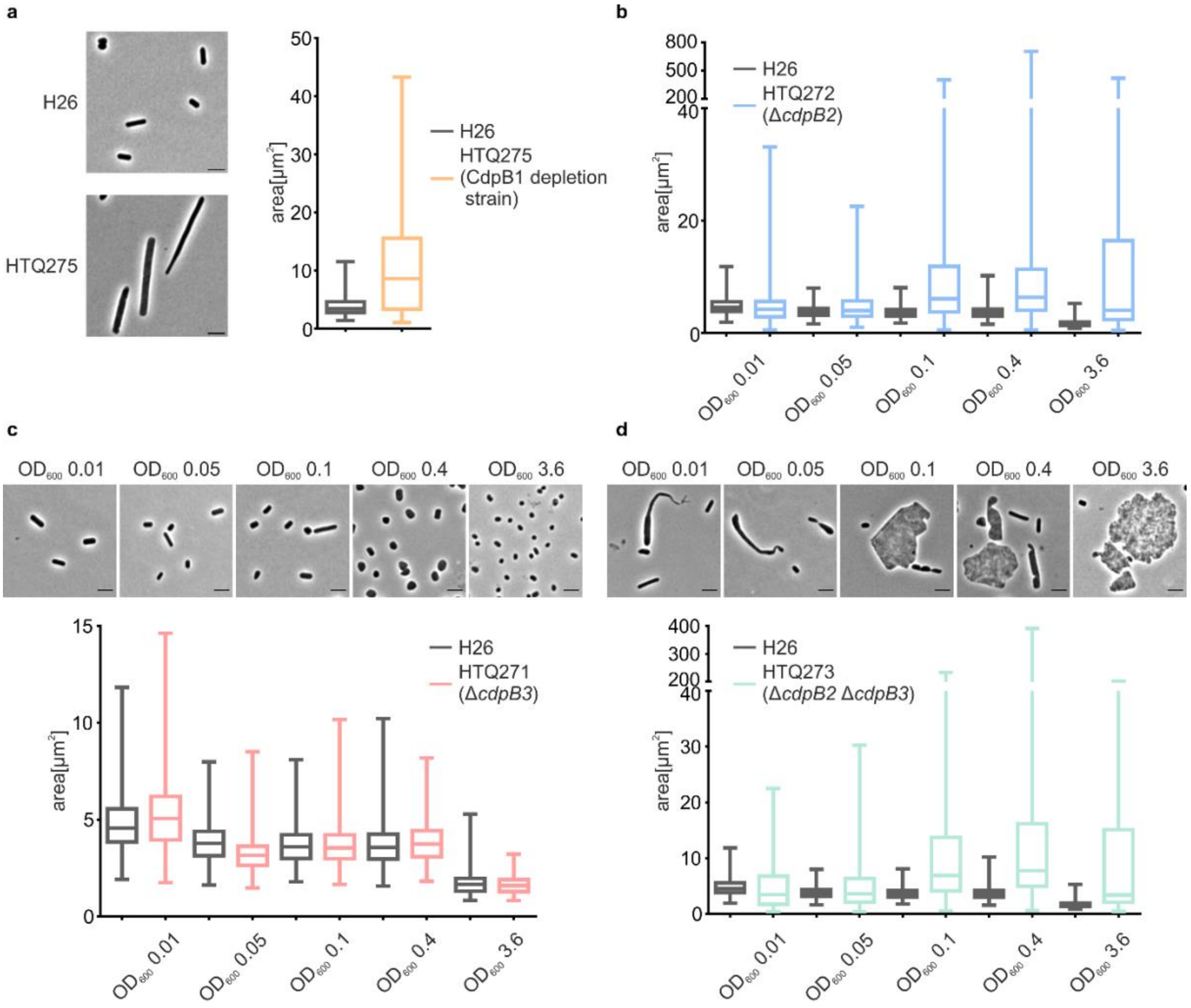


**Supplementary Figure 4: Cell shape analysis of the *cdpB* mutation strains. a** Microscopy images of the wildtype strain H26 and the CdpB1 depletion strain HTQ275 at early exponential phase. The cell shape analysis of H26 (grey) and HTQ275 (orange) from three independent experiments is summarized in the box blots with >1000 cells analyzed per strain. The whiskers indicate the minimum and maximum value and the bar indicates the mean. **b** Cell shape analysis of H26 (grey) and the *cdpB2* (blue) deletion strain HTQ272 at different growth stages (images shown in Fig. 2d). The box blots summarize the result of three independent experiments with >1000 cells per strain at a certain $OD_{600}$. The whiskers indicate the minimum and maximum value and the bar indicates the mean. **c** Microscopy images of the wildtype strain H26 and the *cdpB3* deletion strain HTQ271 at different growth phases. The cell shape analysis of H26 (grey) and HTQ271 (red) from three independent experiments is summarized in the box blots with >1000 cells analyzed per strain and growth stage. The whiskers indicate the minimum and maximum value and the bar indicates the mean of each sample. **d** Microscopy images of the wildtype strain H26 and the *cdpB2cdpB3* double deletion strain HTQ273 at different growth phases. The cell shape analysis of H26 (grey) and HTQ273 (green) from three independent experiments is summarized in the box blots with >1000 cells analyzed per strain and growth stage. The whiskers indicate the minimum and maximum value and the bar indicates the mean of each sample. The mean cell areas of each strain are indicated in Supplementary Table 2.

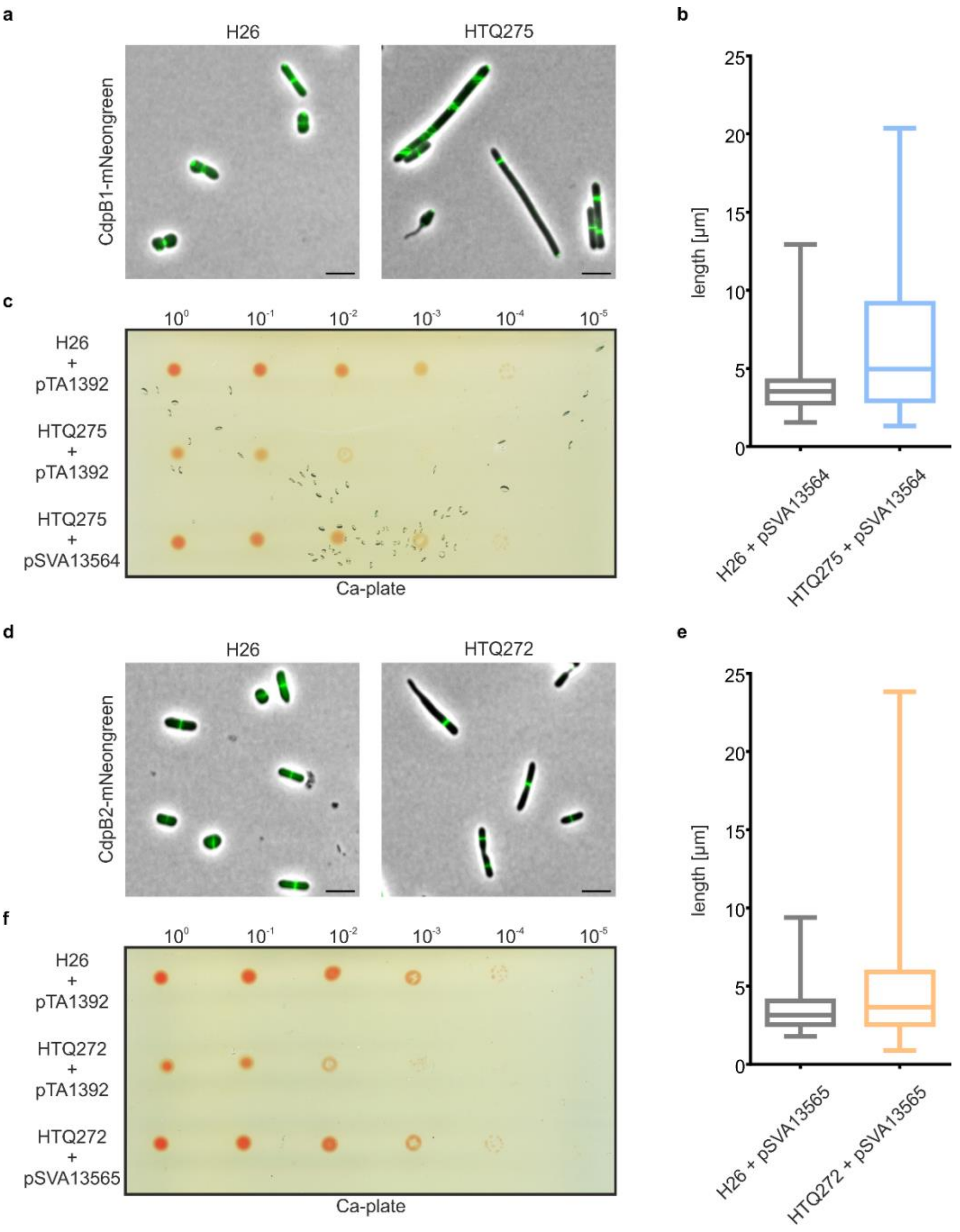


**Supplementary Figure 5: Complementation test of CdpB1- / CdpB2-mNeongreen constructs in the respective mutation strains. a** Fluorescence microscopy of wildtype cells (H26) and cells of the CdpB1 depletion strain (HTQ275) expressing CdpB1-mNeongreen. Scale bar: 4 µm. **b** Cell shape analysis of the imaged cells showing that expression of CdpB1-mNeongreen does not complement the cell shape phenotype of HTQ275. The blot summarizes the result of three independent replicates including > 1000 cells per strain. **c** Spot dilution assay of H26, HTQ275 both transformed with an empty expression plasmid (pTA1392) to complement uracil auxotrophy and the HTQ275 strain expressing CdpB1-mNeongreen. Based on a starting $OD_{600}$ of 0.2 a serial dilution of the strains was made with a final dilution at $10^{-5}$. The diluted cells were spotted on Ca-plates. Strain HTQ275 expressing CdpB1-mNeongreen shows restored viability to wildtype levels compared to HTQ275 without CdpB1-mNeongreen expression. Three independent replicates of the experiment were made with same results. **d** Fluorescence microscopy of wildtype cells (H26) and cells of the CdpB2 deletion strain (HTQ272) expressing

CdpB2-mNeongreen. Scale bar: 4 µm. **e** Cell shape analysis of the imaged cells showing that expression of CdpB2-mNeongreen does not complement the cell shape phenotype of HTQ272. The blot summarizes the result of three independent replicates including > 1000 cells per strain. **f** Spot dilution assay of H26, HTQ272 both transformed with an empty expression plasmid (pTA1392) to complement uracil auxotrophy and the HTQ272 strain expressing CdpB2-mNeongreen. Based on a starting $OD_{600}$ of 0.2 a serial dilution of the strains was made with a final dilution at $10^{-5}$. The diluted cells were spotted on Ca-plates. Strain HTQ272 expressing CdpB2-mNeongreen shows restored viability to wildtype levels compared to HTQ272 without CdpB2-mNeongreen expression. Three independent replicates of the experiment were made with same results.

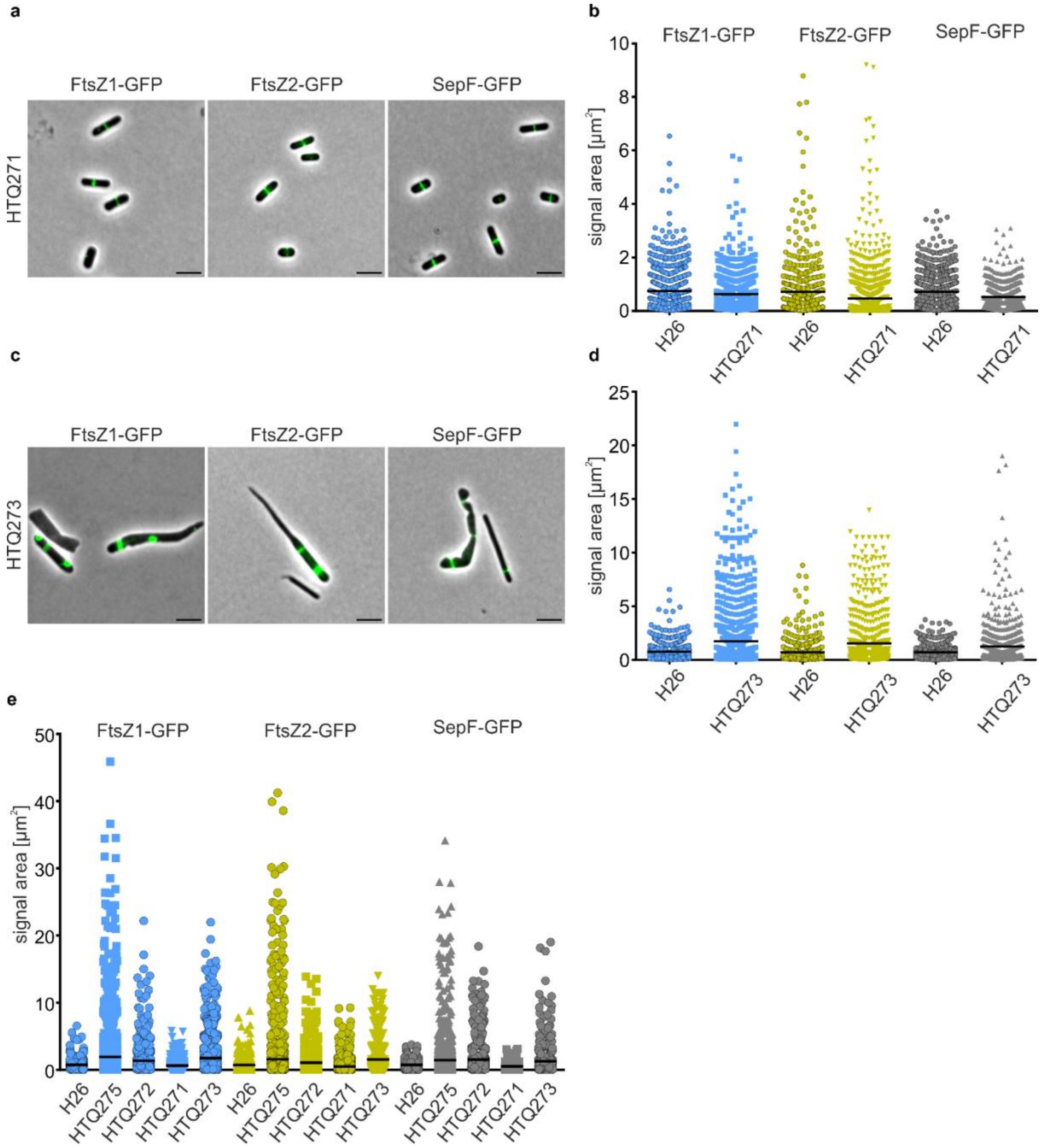


**Supplementary Figure 6: Positioning of cell division proteins of *H volcanii* in Δ*cdpB3* and Δ*cdpB2*Δ*cdpB3* deletion strain. a** Fluorescence microscopy HTQ271 (Δ*cdpB3*) expressing GFP tagged, *ftsZ1*, *ftsZ2* and *sepF*. **b** Analysis of the signal area of each GFP construct in H26 and HTQ271. Each signal area per strain is blotted individually, and the mean is indicated as a black bar. **c** Fluorescence microscopy HTQ273 (Δ*cdpB2*Δ*cdpB3*) expressing GFP tagged, *ftsZ1*, *ftsZ2* and *sepF*. **d** Analysis of the signal area of each GFP construct in H26 and HTQ273. Each signal area per strain is blotted individually, and the mean is indicated as a black bar. The results comprise data from three independent replications per strain and GFP construct with > 1000 cells being analyzed. Scale bar: 4 µm. **e** Summary of the signal areas GFP tagged , *ftsZ1*, *ftsZ2* and *sepF* measured in all strains. The mean signal area of each protein of each strain are indicated in Supplementary Table 1.

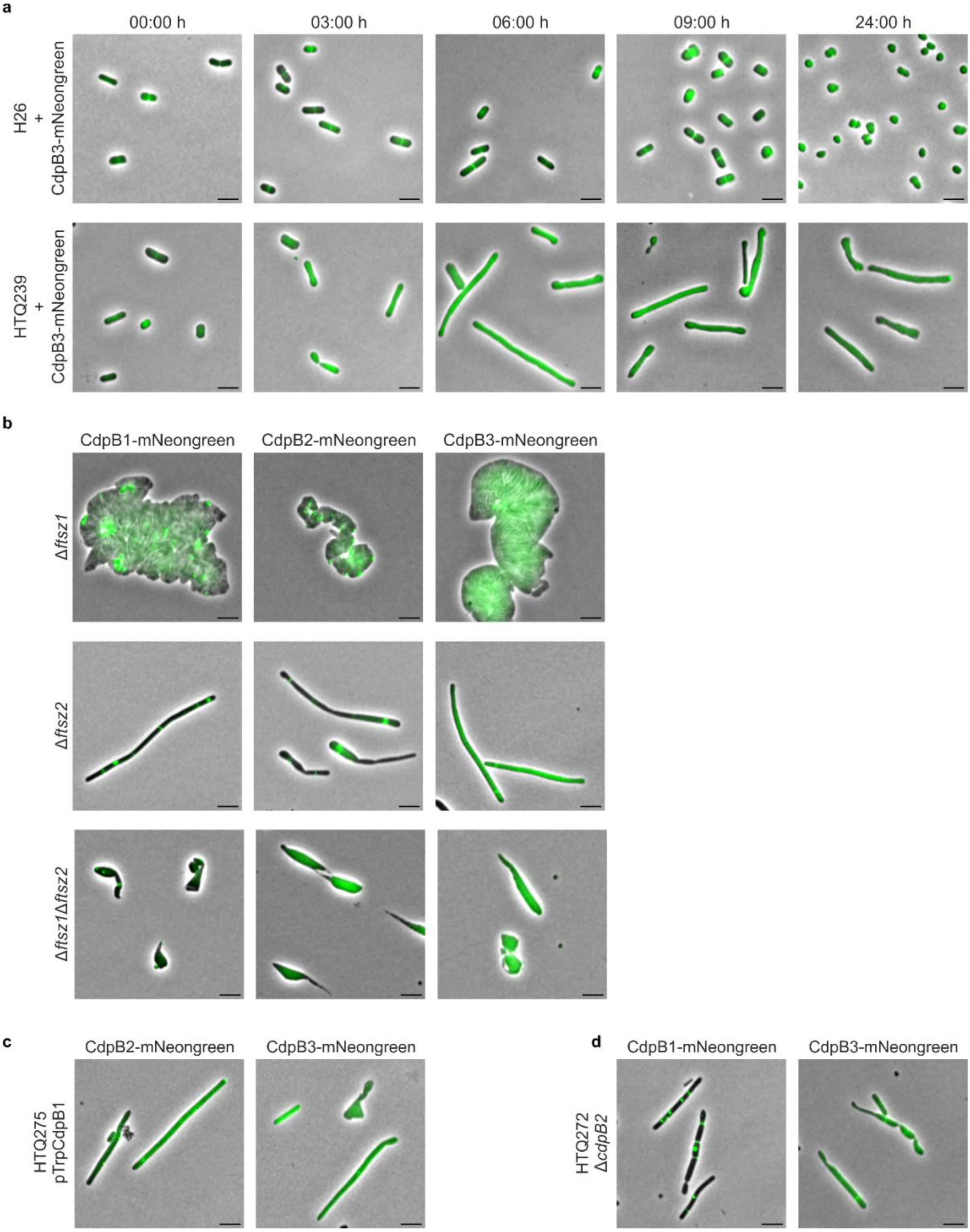


**Supplementary Figure 7: CdpB3 localization during SepF depletion, CdpB1-3 localization in *ftsZ1* and *ftsZ2* deletion strain and localization of the CdpB proteins in absence of CdpB1 or CdpB2. a** Fluorescence microscopy of H26 and the SepF depletion strain HTQ239 expressing CdpB3-mNeongreen from plasmid under the control of its native promotor. Cells were imaged before (00:00 h) and at different timepoints after SepF depletion was induced. **b** Fluorescence microscopy of Δ*ftsZ1,* Δ*ftsZ2* or Δ*ftsZ1*Δ*ftsZ2* strains expressing either CdpB1-, CdpB2 or CdpB3-mNeongreen. **c** Fluorescence microscopy of the CdpB1 depletion strain expressing either CdpB2-mNeongreen or CdpB3-mNeongreen. **d** Fluorescence microscopy of the CdpB2 deletion strain expressing either

CdpB1- mNeongreen or CdpB3-mNeongreen. All experiments were repeated three times with the same result. Scale bar: 4 µm.

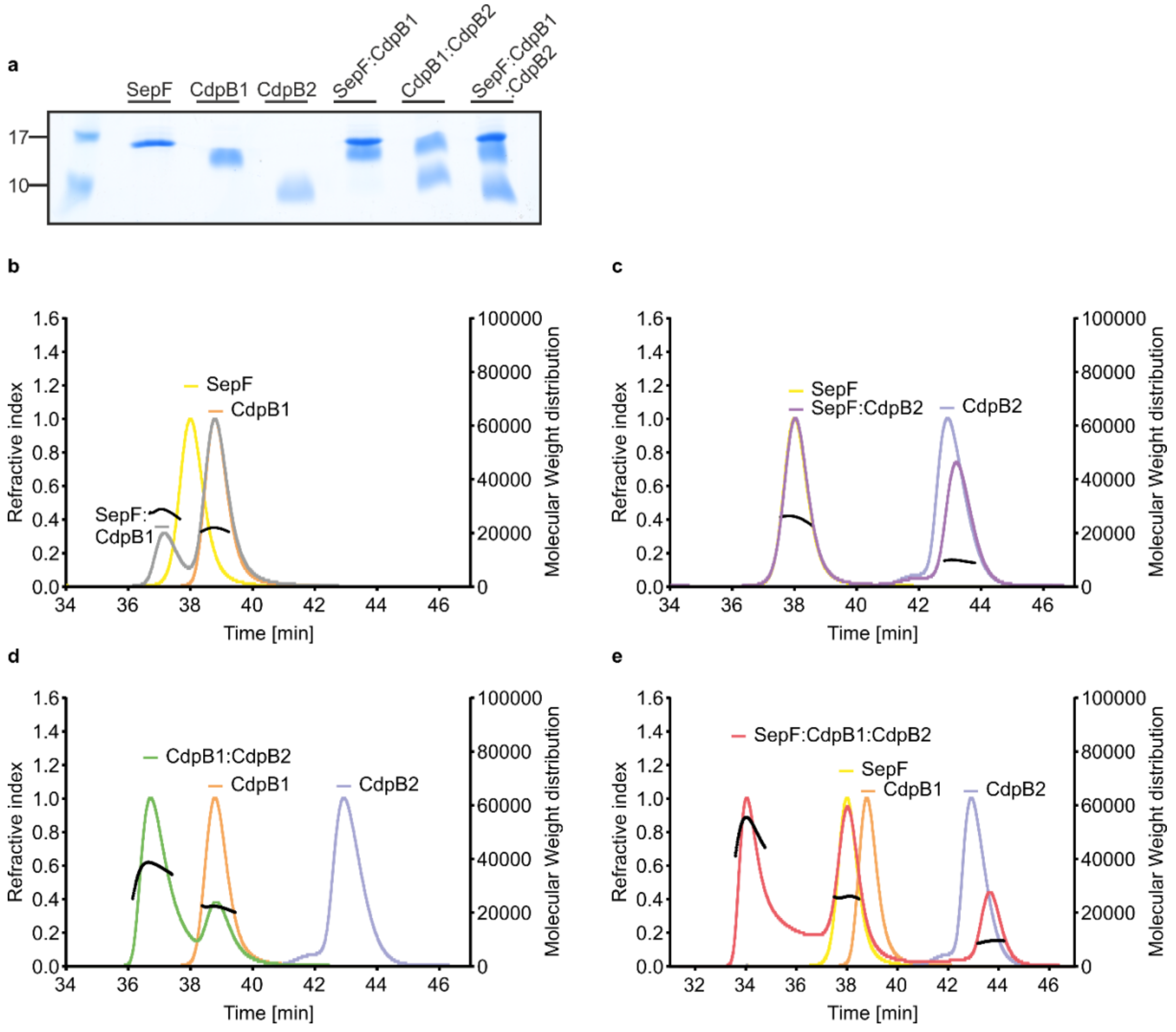


**Supplementary Figure 8: SDS-gel of the peak fractions of the *A. fulgidus* SEC runs and SEC-MALS runs of the *H. volcanii* proteins. a** SDS-gel of the peak elution fractions of the SEC runs of SepF, CdpB1 and CdpB3 alone as well as from the SEC runs of the mixed samples SepF:CdpB1, CdpB1:CdpB2 and CdpB1: CdpB2:SepF (*A. fulgidus proteins*). **b-e** SEC-MALS with proteins from *H. volcanii*: **b** SepF (yellow) and CdpB1 (orange) alone and SepF:CdpB1 together (grey). **c** SepF (yellow) and CdpB2 (blue) alone and SepF:CdpB2 together (purple). **d** CdpB1 (orange) and CdpB2 (blue) alone and CdpB1:CdpB2 together (green). **e** SepF (yellow), CdpB1 (orange) and CdpB2 (blue) alone and SepF:CdpB1:CdpB2 together (red).

**Supplementary Table 1.** Mean signal areas of FtsZ1, FtsZ2 and SepF together with the standard error of the mean measured in the different *H volcanii* strains of this study.

| **Signal area [$\mu m^2$] for** | **FtsZ1** | | **FtsZ2** | | **SepF** | |
|---|---|---|---|---|---|---|
| | Mean | SEM | Mean | SEM | Mean | SEM |
| H26 (WT) | 0.75 | 0.02 | 0.72 | 0.05 | 0.73 | 0.02 |
| HTQ275 (pTrpCdpB1) | 1.92 | 0.09 | 1.60 | 0.1 | 1.50 | 0.06 |
| HTQ272 (Δ*cdpB2*) | 1.35 | 0.08 | 1.10 | 0.05 | 1.50 | 0.07 |
| HTQ271 (Δ*cdpB3*) | 0.62 | 0.02 | 0.50 | 0.02 | 0.5 | 0.02 |
| HTQ273 (Δ*cdpB2*Δ*cdpB3*) | 1.73 | 0.08 | 1.60 | 0.08 | 1.30 | 0.07 |

**Supplementary Table 2**. Mean signal cell area at different optical densities (OD) together with the standard error of the mean measured in the different *H volcanii* strains of this study.

| **Cell area [µm$^2$] at** | **$OD_{600}$ 0.01** | | **$OD_{600}$ 0.05** | | **$OD_{600}$ 0.1** | | **$OD_{600}$ 0.4** | | **$OD_{600}$ 3.6** | |
|---|---|---|---|---|---|---|---|---|---|---|
| | Mean | SEM | Mean | SEM | Mean | SEM | Mean | SEM | Mean | SEM |
| H26 | 4.5 | 0.04 | 3.7 | 0.03 | 3.5 | 0.02 | 3.5 | 0.01 | 1.6 | 0.004 |
| HTQ272 (Δ*cdpB2*) | 4.1 | 0.11 | 4.3 | 0.08 | 10.6 | 0.59 | 15.4 | 0.64 | 10.7 | 0.33 |
| HTQ271 (Δ*cdpB3*) | 4.8 | 0.04 | 3.1 | 0.02 | 3.5 | 0.02 | 3.6 | 0.01 | 1.5 | 0.004 |
| HTQ273 (Δ*cdpB2*Δ*cdpB3*) | 4.1 | 0.08 | 3.8 | 0.06 | 11.9 | 0.58 | 14.8 | 0.49 | 9.7 | 0.29l |

**Supplementary Table 3**. Data collection and refinement statistics (Molecular Replacement).

| | CdpB1 & CdpB22 |
|---|---|
| **Data collection** | |
| Space group | P 21 21 21 |
| Wavelength (Å) | 0.9999 |
| *a*, *b*, *c* (Å) | 71.1, 102.6, 112.3, |
| α, β, γ (°) | 90, 90, 90 |
| Resolution (Å) | 39.02 - 2.29 (2.37 - 2.29) |
| $R_{meas}$ | 0.048 (1.6) |
| CC1/2 | 1.0 (0.5) |
| I/σ(I) | 11.3 (1.5) |
| Completeness (%) | 89 (49) |
| Multiplicity | 6 (6.5) |
| **Refinement** | |
| Number of reflections | 30556 (463) |
| $R_{work}$ / $R_{free}$ | 0.23/ 0.28 |
| Ramachandran favored (outliers) (%) | 96 |
| Clashscore | 8 |
| Rotamer outliers (%) | 1.2 |
| No. atoms | |
| Protein | 5083 |
| Ligand/ion | N/A |
| Water | 4 |
| *B*-factors | |
| Protein | 68 |
| Ligand/ion | N/A |
| Water | 52 |
| R.m.s. deviations | |
| Bond lengths (Å) | 0.008 |
| Bond angles (°) | 1.03 |

The experiment was executed on a single crystal. Statistics for the highest-resolution shell are shown in parentheses.

**Supplementary Table 4**: Strains used in this study

| **Strain Name** | **Background strain** | **Genotype** | **Source/reference** |
|---|---|---|---|
| ***H. volcanii*** | | | |
| H26 | - | *ΔpyrE2* | (1) |
| H98 | - | *ΔpyrE2 ΔhdrB* | (2) |
| HTQ271 | H26 | *ΔpyrE2 ΔcdpB3* | This study |
| HTQ272 | H26 | *ΔpyrE2 ΔcdpB2* | This study |
| HTQ273 | HTQ272 | *ΔpyrE2 ΔcdpB2 ΔcdpB3* | This study |
| HTQ275 | H98 | *ΔpyrE2 ΔhdrB cdpB1*::[*p.tnaA-cdpB1-hdrB*$^{+}$] | This study |
| HTQ239 | H98 | *ΔpyrE2 ΔhdrB hvo_0392*::[*p.tnaA-hvo_0392-hdrB*$^{+}$] | (3) |
| ID76 | H98 | *ΔpyrE2 ΔhdrB p.fdx-hdrB ΔftsZ1* | (4) |
| ID77 | H98 | *ΔpyrE2 ΔhdrB p.fdx-hdrB ΔftsZ2* | (4) |
| ID112 | ID77 | *ΔpyrE2 ΔhdrB p.fdx-hdrB ΔftsZ1 ΔftsZ2* | (4) |
| ***E. coli*** | | | |
| 10-beta Competent Cells "TOP10" | - | *Δ(ara-leu) 7697 araD139 fhuA ΔlacX74 galK16 galE15 e14- φ80*d*lacZΔM15 recA1 relA1 endA1 nupG rpsL* (Str$^{R}$) *rph spoT1 Δ(mrr-hsdRMS-mcrBC)* | New England Biolabs |
| *dam*$^{-}$/*dcm*$^{-}$<br>Competent Cells | - | *ara-14 leuB6 fhuA31 lacY1 tsx78 glnV44 galK2 galT22 mcrA dcm-6 hisG4 rfbD1 R(zgb210::Tn10)* Tet$^{S}$ *endA1 rspL136* (Str$^{R}$) *dam13::Tn9* (Cam$^{R}$) *xylA-5 mtl-1 thi-1 mcrB1 hsdR2* | New England Biolabs |
| Rosetta™(DE3)<br>Competent Cells | - | F$^{-}$ *ompT hsdS*$_{B}$($r_{B}^{-}$ $m_{B}^{-}$) *gal dcm* (DE3) pRARE (Cam$^{R}$) | Novagen |
| BL21 (DE3)<br>Competent Cells | - | F$^{-}$ *ompT hsdS*$_{B}$($r_{B}^{-}$ $m_{B}^{-}$) *gal dcm* (DE3) | Novagen |

**Supplementary Table 5**: Plasmids used in this study

| Plasmids | Description | Primers used | Enzymes used | Source/reference |
|---|---|---|---|---|
| pTA131 | Integrative plasmid with a *pyrE2* selection marker for gene deletions in *H. volcanii* (Amp$^r$) | - | - | (2) |
| pTA1392 | Plasmid for the expression of proteins in *H. volcanii* under control of *p.tnaA* and *pyrE2, hdrB* selection markers (Amp$^r$) | | | (5) |
| pTA1369 | Integrative plasmid with a *pyrE2* and *hdrB* selection marker containing the *p.tnaA1* promotor to generate a tryptophan inducible allele of a gene (Amp$^r$) | - | - | (6) |
| pIDJL-40 | Plasmid for the expression of proteins with a C-terminal smRS-GFP-tag and a *pyrE2, hdrB* selection markers (Amp$^r$) | - | - | (7) |
| pSVA13429 | Plasmid for the expression of proteins with an N-terminal His-SUMO-tag for expression in *E. coli* (Kan$^R$) | | | This study |
| pSVA3943 | Plasmid to express two proteins, one with a c-terminal smRS-GFP tag and on with a c-terminal mCherry tag under the control of *p.tnaA.* *pyrE2* selection marker based on (Amp$^r$) | - | - | (3) |
| pSVA5910 | Plasmid for expression of FtsZ1-GFP under the control of its native promotor (Amp$^r$) | - | - | (3) |
| pSVA5956 | Plasmid for expression of FtsZ2-GFP under the control of its native promotor with a *pyrE2* selection marker (Amp$^r$) | - | - | (3) |
| pSVA5996 | Integrative plasmid for the generation of a *cdpB1* deletion strain (Amp$^r$) | 11086, 12100; 12101, 11089 | BamHI and in vivo ligation | This study |
| pSVA5997 | Plasmid for the expression of proteins with a C-terminal mNeongreen-tag and a *pyrE2, hdrB* selection markers | 11074, 11075 | BamHI, NotI | This study |

| | | | | |
|---|---|---|---|---|
| | (Amp$^r$) | | | |
| pSVA13504 | Plasmid for expression of SepF-GFP under the control of its native promotor with a *pyrE2* selection marker (Amp$^r$) | - | - | (3) |
| pSVA13510 | Integrative plasmid for the exchange of *cdpB1* native promoter with a tryptophan inducible one (Amp$^r$) | 12112, 12113; | NdeI, BamHI; BglII | This study |
| pSVA13521 | Integrative plasmid for the generation of a *cdpB2* deletion strain (Amp$^r$) | 12135, 12136; 12137, 12138 | BamHI and in vivo ligation | This study |
| pSVA13522 | Integrative plasmid for the generation of a *cdpB3* deletion strain (Amp$^r$) | 12139, 12140; 12141, 12142 | BamHI and in vivo ligation | This study |
| pSVA13564 | Plasmid for expression of CdpB1-mNeongreen under the control of its native promotor (Amp$^r$) | 12517, 11085 | ApaI, BamHI | This study |
| pSVA13565 | Plasmid for expression of CdpB2-mNeongreen under the control of its native promotor (Amp$^r$) | 12518, 12121 | ApaI, BamHI | This study |
| pSVA13566 | Plasmid for expression of CdpB3-mNeongreen under the control of its native promotor (Amp$^r$) | 12519, 12115 | ApaI, BamHI | This study |
| pSVA13595 | Tryptophan inducible SepF-GFP, CdpB1-mCherry double expression plasmid with a *pyrE2* selection marker (Amp$^r$) | 12565; 11085<br><br>8096, 8802 | EcoRI, BamHI (for CdpB1)<br><br>NdeI, NheI (for SepF) | This study |
| pSVA13596 | Tryptophan inducible SepF-GFP, CdpB2-mCherry double expression plasmid with a *pyrE2* selection marker (Amp$^r$) | 12566; 12121<br><br>8096, 8802 | EcoRI, BamHI (for CdpB2)<br><br>NdeI, NheI (for SepF) | This study |
| pSVA13597 | Tryptophan inducible SepF-GFP, CdpB3-mCherry double expression plasmid with a *pyrE2* selection marker (Amp$^r$) | 12567; 12115<br><br>8096, 8802 | EcoRI, BamHI (for CdpB3)<br><br>NdeI, NheI (for SepF) | This study |
| pSVA13718 | Tryptophan inducible CdpB2-GFP, CdpB1-mCherry double | 12565; 11085 | EcoRI, BamHI (for CdpB1) | This study |

| | | | | |
|---|---|---|---|---|
| | expression plasmid with a *pyrE2* selection marker (Amp$^r$) | 12120, 13125 | NdeI, NheI (for CdpB2) | |
| pSVA13719 | Tryptophan inducibleCdpB3-GFP, CdpB1-mCherry double expression plasmid with a *pyrE2* selection marker (Amp$^r$) | 12566; 12121<br><br>12114, 13126 | EcoRI, BamHI (for CdpB1)<br><br>NdeI, NheI (for CdpB3) | This study |
| pSVA13720 | Tryptophan inducible CdpB3-GFP, CdpB2-mCherry double expression plasmid with a *pyrE2* selection marker (Amp$^r$) | 12567; 12115<br><br>12114, 13126 | EcoRI, BamHI (for CdpB2)<br><br>NdeI, NheI (for CdpB3) | This study |
| pSVA13585 | Plasmid for the expression of His-SUMO-CdpB1 from *A. fulgidus* in *E. coli* (Kan$^R$) | 12555, 12556 | In vivo ligation | This study |
| pSVA13586 | Plasmid for the expression of His-SUMO-CdpB2 from *A. fulgidus* in *E. coli* (Kan$^R$) | 12557, 12558 | In vivo ligation | This study |
| pSVA13587 | Plasmid for the expression of His-SUMO-SepF from *A. fulgidus* in *E. coli* (Kan$^R$) | 12559, 12560 | In vivo ligation | This study |
| pSVA13700 | Plasmid for the expression of His-SUMO-SepF from *H. volcanii* in *E. coli* (Kan$^R$) | 12568, 12569 | in vivo ligation | This study |
| pSVA13703 | Plasmid for the expression of His-SUMO-CdpB1 from *H. volcanii* in *E. coli* (Kan$^R$) | 12574, 12575 | in vivo ligation | This study |
| pSVA13704 | Plasmid for the expression of His-SUMO-CdpB2 from *H. volcanii* in *E. coli* (Kan$^R$) | 12576, 12577 | in vivo ligation | This study |

**Supplementary Table 6**: Primers used in this study

| Primer Number/Name | Sequence 5'→3' | Description |
|---|---|---|
| 11086 | GGCGAATTGGGTACCGAGGAAATCG CCTACCACAG | Forward primer for the amplification of the up-stream region of *hvo_1691* (*cdpB1*) with 15 complementary bases to linearized pTA131 |
| 12100 | CGCGGATCCATCCGCCAACATGGGA TGCC | Reverse primer for the amplification of the up-stream region of *hvo_1691* (*cdpB1*) with BamHI restriction site |
| 12101 | CGCGGATCCCGTTCTAACTCCCGTTT CC | Forward primer for the amplification of the down-stream region of *hvo_1691* (*cdpB1*) with BamHI restriction site |
| 11089 | GGCGGCCGCTCTAGAGGCGTATCTC TACCCCTTCG | Reverse primer for the amplification of the down-stream region of *hvo_1691* (*cdpB1*) with 15 complementary bases to linearized pTA131 |
| 12143 | TCTTCGGCCTGCTGGACAAC | Forward primer for the screening of successful CdpB1 depletion strain generation |
| 12144 | ACTTCAAGCGCGACACGACG | Reverse primer for the screening of successful CdpB1 depletion strain generation |
| 11074 | CGCGGATCCATGGTCTCGAAGGGCG AGG | Forward primer for the amplification of *mNeongreen* with a BamHI restriction site (cloned in pIDJL-40) |
| 11075 | TTAGCGGCCGCTCACTTGTAGAGCTC GTCCATG | Reverse primer for the amplification of *mNeongreen* with a NotI restriction site (cloned in pIDJL-40) |
| 12112 | ATTGCGCATATGGACGGGACGCCCG AAGAG | Forward primer for the amplification of *cdpB1* with a NdeI restriction site (cloned in pTA1369) |
| 12113 | GCGGGATCCTCACGCGTAGTCCGGG ACGTCGTACGGGTAGCTGCCGGCGA CCAGCGACTCTTCG | Reverse primer for the amplification of *cdpB1-ha* with a BamHI restriction site (cloned in pTA1369) |
| 11066 | TCTAGAGCGGCCGCCAC | Forward primer for the linearization of pTA131 |
| 11067 | GGTACCCAATTCGCCCTATAGTG | Reverse primer for the linearization of pTA131 |
| 11905 | ACCGTCGACCCGACTGGAAAGCG | Forward primer for the linearization of pSVA13429 |
| 11906 | ACCACCGATCTGTTCGCGATGCG | Reverse primer for the linearization of pSVA13429 |
| 12135 | GGCGAATTGGGTACCGCTGGAAGCT GTCGTTCAC | Forward primer for the amplification of the up-stream region of *hvo_1964* (*cdpB2*) with 15 complementary bases to linearized pTA131 |
| 12136 | CGCGGATCCACACACCCCTTCCTCGT C | Reverse primer for the amplification of the up-stream region of *hvo_1964* (*cdpB2*) with BamHI restriction site |

| | | |
|---|---|---|
| 12137 | CGCGGATCCATTCAGGATAGATTCTCAATGCGG | Forward primer for the amplification of the down-stream region of *hvo_1964* (*cdpB2*) with BamHI restriction site |
| 12138 | GGCGGCCGCTCTAGACGGCATCGAACCGCG | Reverse primer for the amplification of the down-stream region of *hvo_1964* (*cdpB2*) with 15 complementary bases to linearized pTA131 |
| 12147 | TCTTGTTCGCGGCCACGATG | Forward primer for the screening of *cdpB2* deletion |
| 12148 | ACAACGCGGTCCAGTTCCTC | Reverse primer for the screening of *cdpB2* deletion |
| 12139 | GGCGAATTGGGTACCTCCGAACTCGCTCGACAG | Forward primer for the amplification of the up-stream region of *hvo_2019* (*cdpB3*) with 15 complementary bases to linearized pTA131 |
| 12140 | CGCGGATCCACCTCCGGTATCGGACG | Reverse primer for the amplification of the up-stream region of *hvo_2019* (*cdpB3*) with BamHI restriction site |
| 12141 | CGCGGATCCCCGCGGCGACTTCG | Forward primer for the amplification of the down-stream region of *hvo_2019* (*cdpB3*) with BamHI restriction site |
| 12142 | GGCGGCCGCTCTAGAGTCTCGCCGGTGCTGTC | Reverse primer for the amplification of the down-stream region of *hvo_2019* (*cdpB3*) with 15 complementary bases to linearized pTA131 |
| 12151 | GAAGCGAGACGACGACTGAC | Forward primer for the screening of *cdpB3* deletion |
| 12152 | GGAAGCGTGACCAGTCGAAC | Reverse primer for the screening of *cdpB3* deletion |
| 12517 | TTCGGGCCCAGCTACGGGGACAGTTCCAG | Forward primer for the amplification of *cdpB1* with its promotor region and an ApaI restriction site (cloned in pSVA5997) |
| 11085 | CGCGGATCCGGCGACCAGCGACTCTTCG | Reverse primer for the amplification of *cdpB1* with BamHI restriction site (cloned in pSVA5997) |
| 12518 | TTCGGGCCCTCTTGTCGAGAAGACTCGTC | Forward primer for the amplification of *cdpB2* with its promotor region and an ApaI restriction site (cloned in pSVA5997) |
| 12121 | CGCGGATCCCCGTTGGACGACGATGTAATCC | Reverse primer for the amplification of *cdpB2* with BamHI restriction site (cloned in pSVA5997) |
| 12519 | TTCGGGCCCCGACTTCGGAGACCAGAATG | Forward primer for the amplification of *cdpB3* with its promotor region and an ApaI restriction site (cloned in pSVA5997) |
| 12115 | CGCGGATCCGTCGGCGGCTTCGAGC | Reverse primer for the amplification of *cdpB3* with BamHI restriction site (cloned in pSVA5997) |
| 8096 | GGAATTCCATATGGGTATCATGAGTAAGATTCTCGG | Forward primer for the amplification of *sepF* with NdeI restriction site |

| | | |
|---|---|---|
| | | (cloned in pSVA3943) |
| 8802 | TCTAGCTAGCGCCGTTGAGCTTCTGC | Reverse primer for the amplification of *sepF* with a NheI restriction site (cloned in pSVA3943) |
| 12565 | CCGGAATTCATGGACGGGACGCCCGAAG | Forward primer for the amplification of *cdpB1* with EcoRI restriction site (cloned in pSVA3943) |
| 12566 | CCGGAATTCATGGCCGACATACTCGCCGAG | Forward primer for the amplification of *cdpB2* with EcoRI restriction site (cloned in pSVA3943) |
| 12567 | CCGGAATTCATGGTCCCCCAGTCACCGACG | Forward primer for the amplification of *cdpB3* with EcoRI restriction site (cloned in pSVA3943) |
| 12120 | ATTGCGCATATGGCCGACATACTCGCCGAGAAC | Forward primer for the amplification of *cdpB2* with NdeI restriction site (cloned in pSVA3943) |
| 13125 | CTAGCTAGCCCGTTGGACGACGATGTAATC | Reverse primer for the amplification of *cdpB2* with a NheI restriction site (cloned in pSVA3943) |
| 12114 | ATTGCGCATATGGTCCCCCAGTCACCGACG | Forward primer for the amplification of *cdpB3* with NdeI restriction site (cloned in pSVA3943) |
| 13126 | CTAGCTAGCGTCGGCGGCTTCGAGC | Reverse primer for the amplification of *cdpB3* with a NheI restriction site (cloned in pSVA3943) |
| 12555 | GCGAACAGATCGGTGGTATGATCGGTGAAATTACCACC | Forward primer for the amplification of codon optimized *cdpB1* from *A. fulgidus* with 17 complementary bases to linearized pSVA13429 |
| 12556 | CCAGTCGGGTCGACGGTTTACTCTTGCCACTCTTCGC | Reverse primer for the amplification of codon optimized *cdpB1* from *A. fulgidus* with 17 complementary bases to linearized pSVA13429 |
| 12557 | GCGAACAGATCGGTGGTATGTATGTGCCGGCGCGTAG | Forward primer for the amplification of codon optimized *cdpB2* from *A. fulgidus* with 17 complementary bases to linearized pSVA13429 |
| 12558 | CCAGTCGGGTCGACGGTTTAACGAACACGACGACGATC | Reverse primer for the amplification of codon optimized *cdpB2* from *A. fulgidus* with 17 complementary bases to linearized pSVA13429 |
| 12559 | GCGAACAGATCGGTGGTATGGGTATCATTGATAAGATCCTGG | Forward primer for the amplification of codon optimized *sepF* from *A. fulgidus* with 17 complementary bases to linearized pSVA13429 |
| 12560 | CCAGTCGGGTCGACGGTTTAACGGCTGCTGCTACG | Reverse primer for the amplification of codon optimized *sepF* from *A. fulgidus* with 17 complementary bases to linearized pSVA13429 |
| 12568 | GCGAACAGATCGGTGGTATGGGTATCATGAGTAAGATTC | Forward primer for the amplification of *sepF* from *H. volcanii* with 17 |

| | | |
|---|---|---|
| | | complementary bases to linearized pSVA13429 |
| 12569 | CCAGTCGGGTCGACGGTTCAGCC GTTGAGCTTC | Reverse primer for the amplification of *sepF* from *H. volcanii* with 17 complementary bases to linearized pSVA13429 |
| 12574 | GCGAACAGATCGGTGGTATGGAC GGGACGCCCGAAG | Forward primer for the amplification of *cdpB1* from *H. volcanii* with 17 complementary bases to linearized pSVA13429 |
| 12575 | CCAGTCGGGTCGACGGTTCAGGC GACCAGCGACTCTTC | Reverse primer for the amplification of *cdpB1* from *H. volcanii* with 17 complementary bases to linearized pSVA13429 |
| 12576 | GCGAACAGATCGGTGGTATGGCC GACATACTCGCCGAG | Forward primer for the amplification of *cdpB2* from *H. volcanii* with 17 complementary bases to linearized pSVA13429 |
| 12577 | CCAGTCGGGTCGACGGTCTACCG TTGGACGACGATGTAATC | Reverse primer for the amplification of *cdpB2* from *H. volcanii* with 17 complementary bases to linearized pSVA13429 |

**References**


1. Bitan-Banin,G., Ortenberg,R. and Mevarech,M. (2003) Development of a Gene Knockout System for the Halophilic Archaeon Haloferax volcanii by Use of the pyrE Gene. *J. Bacteriol.*, **185**, 772–778.

2. Allers,T., Ngo,H.-P., Mevarech,M. and Lloyd,R.G. (2004) Development of additional selectable markers for the halophilic archaeon *Haloferax volcanii* based on the *leuB* and *trpA* genes. *Appl. Environ. Microbiol.*, **70**, 943–53.

3. Nußbaum,P., Gerstner,M., Dingethal,M., Erb,C. and Albers,S.V. (2021) The archaeal protein SepF is essential for cell division in Haloferax volcanii. *Nat. Commun.*, **12**, 1–15.

4. Liao,Y., Ithurbide,S., Evenhuis,C., Löwe,J. and Duggin,I.G. (2021) Cell division in the archaeon Haloferax volcanii relies on two FtsZ proteins with distinct functions in division ring assembly and constriction. *Nat. Microbiol.*, **6**, 594–605.

5. Gamble-Milner,R. (2016) Genetic analysis of the Hel308 helicase in the archaeon Haloferax volcanii. *PhD thesis, Univ. Nottingham*.

6. Braun,F., Thomalla,L., van der Does,C., Quax,T.E.F., Allers,T., Kaever,V. and Albers,S.-V. (2019) Cyclic nucleotides in archaea: Cyclic di-AMP in the archaeon Haloferax volcanii and its putative role. *Microbiologyopen*, **8**, 1–23.

7. Duggin,I.G., Aylett,C.H.S., Walsh,J.C., Michie,K.A., Wang,Q., Turnbull,L., Dawson,E.M., Harry,E.J., Whitchurch,C.B., Amos,L.A., *et al.* (2015) CetZ tubulin-like proteins control archaeal cell shape. *Nature*, **519**, 362–365.
